# Fine-scale ecological and transcriptomic data reveal niche differentiation of an allopolyploid from diploid parents in *Cardamine*

**DOI:** 10.1101/600783

**Authors:** Reiko Akiyama, Jianqiang Sun, Masaomi Hatakeyama, Heidi E.L. Lischer, Roman V. Briskine, Angela Hay, Xiangchao Gan, Miltos Tsiantis, Hiroshi Kudoh, Masahiro M. Kanaoka, Jun Sese, Kentaro K. Shimizu, Rie Shimizu-Inatsugi

## Abstract

Polyploidization, or whole genome duplication, is one of the major mechanisms of plant speciation. Allopolyploids (species that harbor polyploid genomes originating from hybridization of different diploid species) have been hypothesized to occupy a niche with intermediate, broader, or fluctuating environmental conditions compared with parental diploids. It remains unclear whether empirical data support this hypothesis and whether specialization of expression patterns of the homeologs (paralogous gene copies resulting from allopolyploidization) relates to habitat environments. Here, we studied the ecology and transcriptomics of a wild allopolyploid *Cardamine flexuosa* and its diploid parents *C. hirsuta* and *C. amara* at a fine geographical scale in their native area in Switzerland. We found that the diploid parents favored opposite extremes in terms of soil moisture, soil carbon-to-nitrogen ratios, and light availability. The habitat of the allopolyploid *C. flexuosa* was broader compared with those of its parental species and overlapped with those of the parents, but not at its extremes. In *C. flexuosa*, the genes related to water availability were overrepresented among those at both the expression level and the expression ratio of homeolog pairs, which varied among habitat environments. These findings provide empirical evidence for niche differentiation between an allopolyploid and its diploid parents at a fine scale, where both ecological and transcriptomic data indicated water availability to be the key environmental factor for niche differentiation.

**Significance statement:** Polyploidization, or whole genome duplication, is common in plants and may contribute to their ecological diversification. However, little is known about the niche differentiation of wild allopolyploids relative to their diploid parents and the gene expression patterns that may underlie such ecological divergence. We detected niche differentiation between the allopolyploid *Cardamine flexuosa* and its diploid parents *C. amara* and *C. hirsuta* along water availability gradient at a fine scale. The ecological differentiation was mirrored by the dynamic control of water availability-related gene expression patterns according to habitat environments. Thus, both ecological and transcriptomic data revealed niche differentiation between an allopolyploid species and its diploid parents.

---

Polyploidization is pervasive in plants (1) and is one of the major mechanisms of speciation (2, 3). In allopolyploids (species with multiple genome sets from different diploid parents), differential expression of homeologs (potentially redundant copies of genes derived from the parent species) has been speculated to reflect selective use of the homeologs in responding to various environmental conditions (4, 5) in addition to higher evolvability due to redundancy (6, 7). This, in turn, might enable allopolyploids to thrive in a range of environments. Consistent with this idea, manipulative experiments in laboratory conditions demonstrated homeolog expression ratio changes in response to environmental factors such as temperature, water availability, and metal concentration (8–12, but see 13). Recent empirical studies show that a small proportion (ca. 1%) of all genes exhibit a ratio change, many of which are known to be involved in response to the specific conditions that were manipulated in the experiment (10, 12). These results suggest that allopolyploids selectively express the homeologs relevant for an appropriate response to the environment. Although the expression patterns in the laboratory and in naturally fluctuating environments, or *in natura*, may be distinct (14–16), little is known about environment-dependent changes in the homeolog expression ratio of allopolyploids in natural habitats and about the divergence of expression patterns from progenitors.

By contrast with the general assumption that the niches of allopolyploids would be distinct, broader, or intermediate compared with that of the parents’, some studies reported that allopolyploid niches are not necessarily intermediate or broad (17–22). The diverse patterns of niche differentiation of allopolyploids from the parents can be attributed in part to the spatial scale of a study. Ecological niche modeling approaches adopted in most studies evaluate environmental factors at a resolution of >1 km^2^, without addressing the possibility that environmental factors varying at smaller spatial scales are associated with niche differentiation (19, 22). If this is the case, fine-scale empirical studies would have a better ability to detect niche differentiation because they can match the scale of species distribution with associated environmental factors (23). Another infrequently considered aspect of niche differentiation studies is environmental fluctuation. Data from multiple time points should enable the evaluation of temporal variation in niche environments as well as differential gene expression (24–27).

The genus *Cardamine* of Brassicaceae has long been a model to study ecological polyploid speciation (28–30). The allotetraploid *Cardamine flexuosa* in Europe derived from diploid parents *C. amara* (genome size 217–273 Mb) and *C. hirsuta* (genome size 198–225 Mb) (31–34, in-house measurement) offers a promising system with which to study the niche differentiation and associated homeolog expression of allopolyploids in comparison with the diploid parents. *Cardamine flexuosa* is estimated to have emerged between 10,000 and one million years ago (33). Although the distribution of the three species largely overlap at the scale of >5 km^2^ (35), anecdotal reports suggest niche differentiation among species, as *C. hirsuta* plants are typically observed on roadsides and in ditches, *C. amara* on river banks and in wet woodlands, and *C. flexuosa* along forest roads (36–42). In a manipulative laboratory experiment where the three species underwent drought, submergence, or fluctuation of the two, *C. hirsuta* performed the best in drought and worst in submergence, the opposite was the case for *C. amara*, and *C. flexuosa* performed similarly well in all treatments (28). The three species thus appear to show niche differentiation along hydrological gradients, characterized by two distinct stresses—water-logging and drought—at the two ends of the gradient at a scale smaller than 5 km^2^. Microarray analyses from the laboratory experiment showed that the gene expression pattern of *C. flexuosa* was similar to that of *C. hirsuta* under drought and to that of *C. amara* under submergence, and that the induction level of most genes in response to water stress was intermediate in *C. flexuosa* compared with that of parental species (28). These results suggested that *C. flexuosa* became a generalist by the transcriptomic plasticity combining two parents, but the response was attenuated as a trade-off. It is yet to be shown whether this applies to the field conditions. In addition, a major limitation of the usage of the *Arabidopsis* microarray was that it allowed the quantification of only the sum of the expression levels of two homeologs of only about 46% of the entire genes. Recently, a tool for analyzing the homeolog expression ratio change in allopolyploids (10) and the genome sequence of *C. hirsuta* (34) became available, enabling evaluation of the expression of homeologs across genomes of *C. flexuosa*. These technical advances and findings from the laboratory experiment make *C. flexuosa* one of few allopolyploids with distinct ecologies and genomic tools (43); however, quantitative data from natural habitats are still lacking.

Here, we conducted a fine-scale study over two seasons to quantify water availability, soil properties, and light availability in the habitats of *C. flexuosa, C. amara*, and *C. hirsuta* within their native range in Switzerland and to analyze homeolog expression patterns of *C. flexuosa* in comparison with parents. Based on previous studies, we hypothesized that *C. flexuosa* would inhabit a wide water-availability gradient, including sites with fluctuating water availability, and that it would differentially express homeologs similarly to either of the parents depending on habitat environments. We addressed the following questions: (1) What is the relative contribution of different environmental factors to the distribution of the allopolyploid *C. flexuosa* as well as the two diploid parental species? (2) What are the key environmental factors associated with species performance? Do they include water availability? (3) Along the gradients of environmental factors contributing to niche differentiation, is *C. flexuosa* in a broader range compared with *C. amara* and *C. hirsuta*? (4) How does the gene expression of *C. flexuosa* relate to that of the diploid parents? (5) Do the ecological and transcriptomic data indicate the same key environmental factor?

## Results

### Habitat Environment of the Allopolyploid *Cardamine flexuosa* and its Diploid Parents

To characterize the habitat environment of the three species in this study, we measured soil properties and light availability at 17 sites in three areas (IR, WBH, and KT) in Switzerland during two seasons (Figs S1–S3, Table S1). Upon these data, the relative contributions of soil properties and light availability to habitat environment were evaluated by principal component analysis (PCA), which suggested three differences among the species. First, sites of two parental diploid species, *C. hirsuta* (red border encompassing red and orange points) and *C. amara* (blue border encompassing black and blue points), were separated (Fig. 1). The separation along the PC1 axis indicates that *C. amara* sites exhibited high soil water content and soil carbon-to-nitrogen (C/N) ratios (Figs 1 and S2*A, B*) as well as a low degree of sky openness (Figs 1 and S3) by contrast with *C. hirsuta* sites. The high C/N ratios in *C. amara* sites likely reflect low nitrogen concentrations in fresh and incubated soils (Figs 1 and S2*C*-*F*). Second, on the PCA plot, sites of the allopolyploid *C. flexuosa* (green border encompassing orange, green, and blue points) overlapped with both parental diploids reflecting the coexisting sites. Importantly, the sites cohabited by *C. flexuosa* and *C. hirsuta* (orange dots) tend to be in the middle of the PC1 axis compared with sites with *C. hirsuta* alone (red dots). A similar tendency, although weaker, was found with *C. amara* (more pronounced in properties such as soil water content, see below and Fig. S2*A*). These analyses suggest that *C. flexuosa* does not occur in extreme habitats compared with the diploid parents. Third, compared with diploid parents, *C. flexuosa* sites were more scattered. This was supported by larger root-mean-square error, which shows the average variance over samples, calculated as square root of the sum of squares of the difference between PC1 value and mean divided by the sample size: *C. flexuosa*: 0.68 and 0.54 in 2013 and 2014, respectively; *C. hirsuta*: 0.48; *C. amara*: 0.27; the same value for both years for diploid parents due to consistency in the sites of occurrence.

**Fig. 1.**
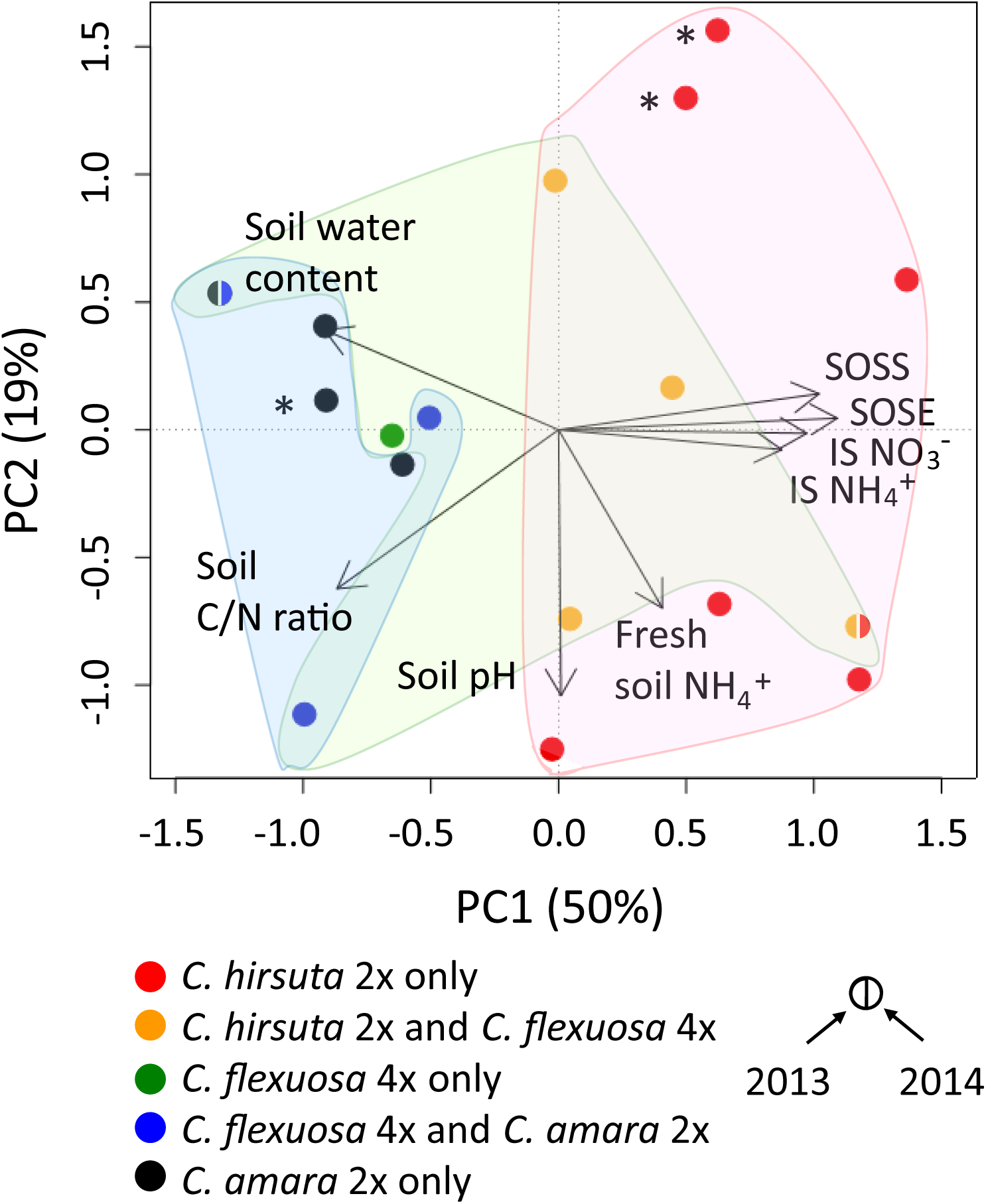
The results of PCA of environmental factors from 17 sites that were subject to the analysis. The relative impacts of different factors are illustrated as vectors. SOSS: Sky openness at season start, SOSE: Sky openness at season end, IS NO_3_^−^: Incubated soil NO_3_^−^, IS NH_4_^+^: Incubated soil NH_4_^+^. The first two PC axes accounted for 50% and 19% of total variation. Different colors of the points indicate different species composition: red, *C. hirsuta* only; orange, *C. hirsuta* and *C. flexuosa*; green, *C. flexuosa* only; blue, *C. flexuosa* and *C. amara*; and black, *C. amara* only. Blue, green, and red background colors indicate the sites with *C. amara, C. flexuosa*, and *C. hirsuta*, respectively. The color of the point is halved for sites with species composition being different between 2013 and 2014. Three sites in the WBH area are indicated with asterisks.

Three sites in the area of WBH (indicated with asterisks in Fig. 1) exemplify the importance of the fine-scale environments. One was occupied by only *C. amara* (black point, WBH1) and the other two by only *C. hirsuta* (red points, WBH2 and 3). This separation would not have been detected at a resolution of >1 km^2^ (Fig. S1*A* and *C*).

By examining each environmental factor separately, we found that the extent of overlap of the habitat of *C. flexuosa* and its parents varied among factors (Figs S2 and S3). *Cardamine flexuosa* was found in an intermediate value range, e.g. for soil water content (*C. amara* 30– 60%, *C. flexuosa* 20–40%, *C. hirsuta* 20–35%, Fig. S2*A*) and sky openness (*C. amara* 5–35%, *C. flexuosa* 5–40%, *C. hirsuta* 10–70%, Fig. S3), all of which contributed to PC1. Therefore, soil water content, soil C/N ratio, and light availability seemed to correlate with habitat differentiation among species, and the habitat environment of *C. flexuosa* generally overlapped with that of the two parental species in the intermediate value range.

### Relative Contributions of Environmental Factors to the Occurrence of the Study Species

To quantify how environmental factors might contribute to the likelihood of occurrence (presence or absence in a given site) of the three species, we performed logistic regression analyses concerning all combinations of environmental factors for each species in each year. In all species, the best model with the lowest Akaike information criterion included water content, C/N ratio, and sky openness at season start. Importantly, the absolute value of the regression coefficient was the largest for water content (Table 1). These results indicate that the three species share environmental factors that determine habitat differentiation and soil water content is the most influential determinant.

**Table 1.** Effects of the environmental factors in the best model on the occurrence of *Cardamine amara, C. hirsuta*, and *C*.*flexuosa* in 2013 and 2014, analysed with logistic multiple regression (*N* = 17). The site of occurrence was consistent in the two years for *C. amara* and for *C. hirsuta*. As *C. flexuosa* occurred in only one of the two years in some sites, we ran four models with different response variables; occurrence in 2013, occurrence in 2014, occurrence in both 2013 and 2014, and occurrence in either 2013 or 2014. For each model, Akaike information criterion (AIC), partial regression coefficient (*β*) ± SE for each variable, and chi-square value (*χ*^2^) and significance (*) are given. * p < 0.05, ** p < 0.01, *** p < 0.001

| Species<br>(ploidy level) | Year | AIC | Water content |  |  | CN ratio |  |  | Sky openness at season start |  |  |
| --- | --- | --- | --- | --- | --- | --- | --- | --- | --- | --- | --- |
| | | | $\beta \pm \text{SE}$ | $\chi^2$ | | $\beta \pm \text{SE}$ | $\chi^2$ | | $\beta \pm \text{SE}$ | $\chi^2$ | |
| *C. amara* (2x) | 2013, 2014 | -2.35 | $0.11 \pm 0.11$ | 8.12 | ** | $0.06 \pm 0.05$ | 7.54 | ** | $-0.01 \pm 0.06$ | 5.69 | * |
| *C. hirsuta* (2x) | 2013, 2014 | -1.99 | $-0.10 \pm 0.10$ | 7.32 | ** | $-0.04 \pm 0.05$ | 6.87 | ** | $0.03 \pm 0.06$ | 7.02 | ** |
| *C. flexuosa* (4x) | 2013 | 6.89 | $-0.08 \pm 0.08$ | 6.81 | ** | $-0.002 \pm 0.04$ | 6.26 | * | $-0.07 \pm 0.05$ | 9.71 | ** |
| | 2014 | 4.47 | $0.03 \pm 0.06$ | 5.47 | * | $-0.02 \pm 0.04$ | 6.31 | * | $-0.08 \pm 0.05$ | 11.33 | *** |
| | 2013 and 2014 | 3.24 | $-0.06 \pm 0.07$ | 5.63 | * | $-0.02 \pm 0.45$ | 6.01 | * | $-0.10 \pm 0.06$ | 13.38 | *** |
| | 2013 or 2014 | 8.07 | $0.003 \pm 0.06$ | 5.59 | * | $-0.001 \pm 0.04$ | 6.36 | * | $-0.05 \pm 0.04$ | 8.43 | ** |

\* $p < 0.05$ , \*\* $p < 0.01$ , \*\*\* $p < 0.001$

The directions of the regression coefficients for the three factors differed among species (Table 1). The occurrence of *C. amara* was associated positively with water content and C/N ratio, and negatively with sky openness at the season’s start. By contrast, the occurrence of *C. hirsuta* was associated with the three factors in the opposite direction. In *C. flexuosa*, the directions of these regression coefficients consisted of a combination of those of its diploid parents. For water content, negative and positive values were observed in 2013 and 2014, respectively, coinciding with the exclusion and inclusion of sites with extreme soil moisture (filled and open arrows in Fig. 2*B* and *F*, Table 1). Furthermore, the values of the regression coefficients for water content and C/N ratio of *C. flexuosa* were generally intermediate between those for *C. amara* and *C. hirsuta*. Hence, these quantitative analyses support the difference of the three species in the PCA plot (Fig. 1), in which the habitat environment of the two diploid parents are distinct, and that of the allopolyploid *C. flexuosa* partly overlapped with those of its two parents at the intermediate range.

**Fig. 2.**
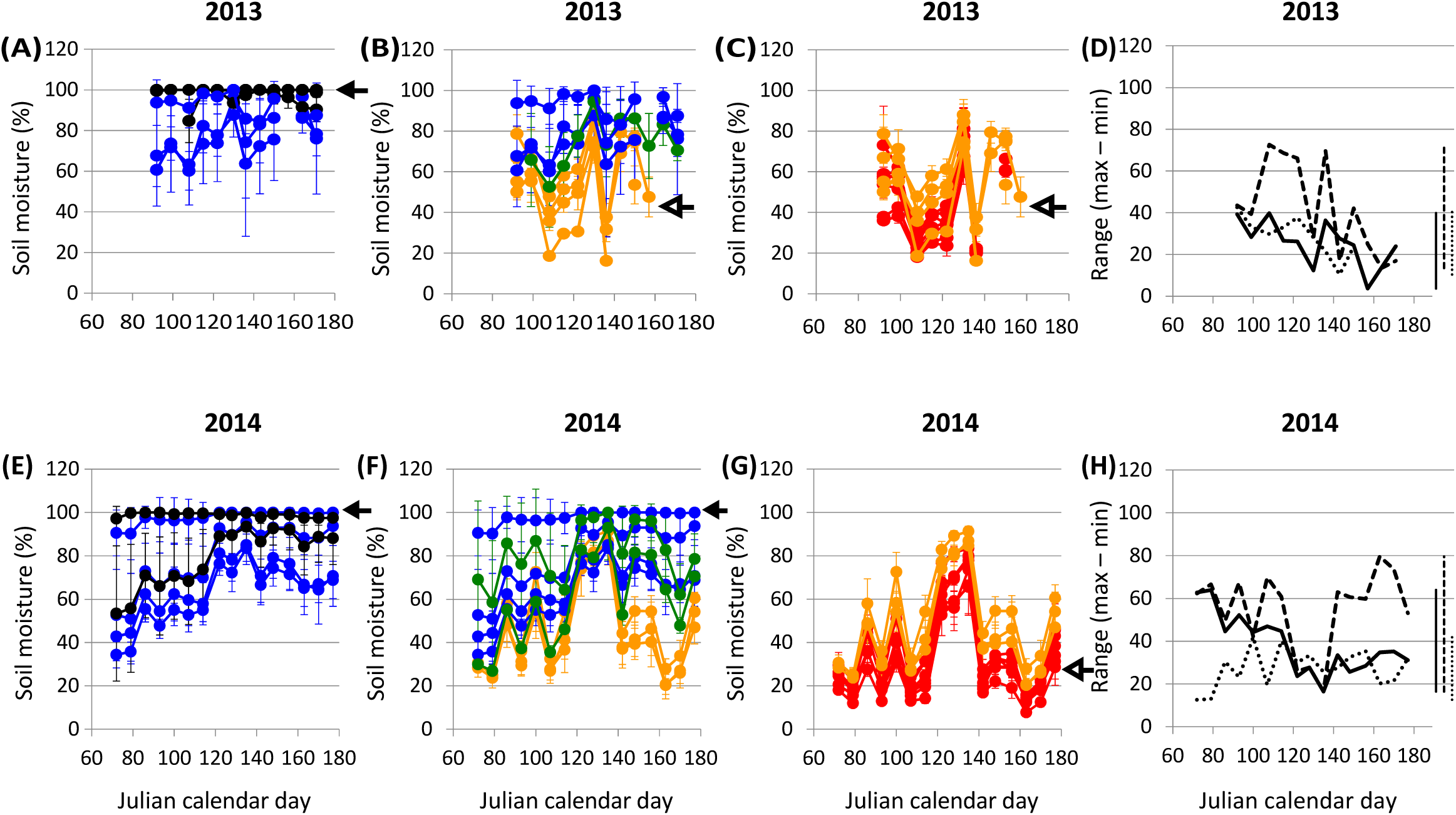
The mean ± standard deviation and the range (maximum–minimum) of soil moisture during growing seasons in 2013 (*A–D*) and 2014 (*E–H*) measured with a soil moisture sensor at each site (3 ≤ *N* ≤ 8 per site). (*A* and *E*) mean soil moisture of sites with *C. amara*. (*B* and *F*) mean soil moisture of sites with *C. flexuosa*. (*C* and *G*) mean soil moisture of sites with *C. hirsuta*. In (*A–C*) and (*E–G*), the color coding of points corresponds to that in Fig. 1 and Fig. S1 to S3, and open and filled arrows indicate the IR5 and KT1 sites, respectively. (*D* and *H*) the range of soil moisture on census dates and for the entire growing season (vertical bars on the right) for each species; solid, dotted, and finely dotted lines indicate *C. amara, C. flexuosa*, and *C. hirsuta*, respectively. In 2013, *C. flexuosa* did not occur in the wettest site, KT1, inhabited by *C. amara* (Fig. 2*A*, filled arrowhead) but appeared in the driest site, IR5, inhabited by *C. hirsuta* (Fig. 2*C–D*, open arrowheads). The opposite was the case in 2014 (Fig. 2*E–G*).

### Relationship between Environmental Factors and Performance

To evaluate how well the species performs in its habitat, we examined the relationship between fitness components and water content and C/N ratio measured in 2013. The analysis was done only for *C. hirsuta* because the sample sizes for the other species became too small after some plants had been lost to flooding. The regression showed that the seed production of *C. hirsuta* was negatively related to soil water content and C/N ratio when controlling for variation in plant size by including it in the model (Table S2). This result indicates that *C. hirsuta* performed better in dry and nutrient-rich environments.

### Temporal Fluctuation of the Soil Moisture

Next we examined whether temporal patterns of habitat water availability varied among species because the key environmental factor for determining niche differentiation turned out to be water content, which likely fluctuated naturally, and because previous experiments showed that *C. flexuosa* performed similarly well with the submerging treatment, drought treatment, and the alternation of the two (28). To this end, we weekly monitored soil moisture during two consecutive growing seasons using an electrical sensor that allowed nondestructive data collection instead of sampling soil at every census. In 2013, the habitat’s level of soil moisture for the diploid parent *C. amara* was high, by contrast with the other diploid parent *C. hirsuta* (Fig. 2*A* and *C*). At any given time, the level of soil moisture of sites inhabited by only *C. amara* (black points in the figure) and by only *C. hirsuta* (red points in the figure) did not overlap with each other. The habitat’s level of soil moisture for the allopolyploid *C. flexuosa* overlapped partly with either of the parental diploids in the intermediate range, but not at the high or low ends (Fig. 2*B*). The level of soil moisture in 2014 was generally similar to that in 2013, although at the beginning of the season, the levels of soil moisture of the *C. amara* and *C. flexuosa* habitats were lower than they were in 2013 (Fig. 2*A, B, E*, and *F*). Furthermore, the soil moisture range throughout the season was the largest for *C. flexuosa* in both 2013 and 2014 (Fig. 2*D* and *H*). Thus, the habitats of *C. flexuosa* had a broad range of soil moisture overlapping with two diploid parents at the intermediate range, suggesting that *C. flexuosa* are tolerant to both submerging and drought stresses as in the laboratory experiments, but less so than each of the diploid parents in extreme environments. The observed niche differentiation along soil moisture provided a basis for studying homeolog expression patterns of *C. flexuosa* in comparison with its diploid parents along a natural environmental gradient.

### PCA of Homeolog Expression of *C. flexuosa* and its Diploid Parents

To compare the gene expression pattern in the allopolyploid and diploids, we analyzed RNA-sequencing (seq) data of *C. flexuosa* at three times (April 18, May 2, and May 16 in 2013) at three sites with contrasting soil moisture levels (IR1, KT2, and KT5, Fig. 3*A*, Tables S3 and S4). In addition, we analyzed the data of the cohabiting parents, *C. hirsuta* at IR1 and *C. amara* at KT5, on May 2. The tissue samples for all the three species were available only for this date due to nonsynchronizing phenology among species and the loss of individuals of *C. amara* to flooding after May 2 (Tables S3 and S4, see Supporting Information, hereafter SI). We obtained an adequate number of RNA-seq reads for all samples (5–10 million) except for one *C. flexuosa* sample (April 18, IR1) that was excluded from analyses. The number of reads classified into each subgenome was almost equal in all *C. flexuosa* samples (SI and Fig. S4), indicating no clear expression bias at subgenomic level.

**Fig. 3.**
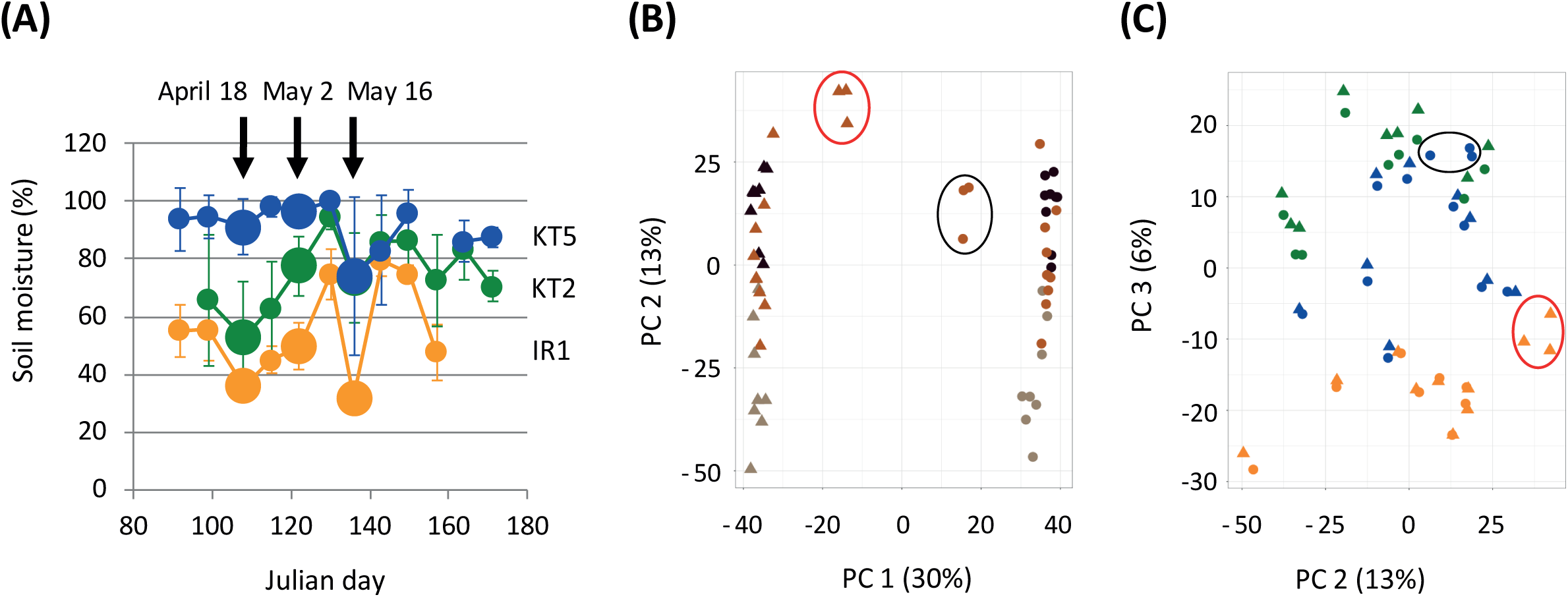
The overview of transcriptomics of the allopolyploid *C. flexuosa* and parents. (*A*) Soil moisture content of the RNA sampling sites in 2013, extracted from Fig. 2*B*. Enlarged points indicate the date of leaf tissue sampling for RNA-seq at IR1 (orange), KT2 (green), and KT5 (blue). *(B)* and *(C)* The plot of PC1 and PC2 (*B*) and PC2 and PC3 (*C*) of the result of PCA of 23,182 homeolog or gene expression in *C. flexuosa* and diploids. Points indicate *C. amara* subgenome (F^a^) and *C. amara* (A, circled in black) while triangles indicate *C. hirsuta* subgenome (F^h^) and *C. hirsuta* (H, circled in red). Different colors in (*B*) indicate different dates: light brown, April 18; brown, May 2; dark brown, May 16. The colors in (*C*) corresponds to the sites as in *(A)* and other figures.

We analyzed the expression pattern of 23,182 homeolog pairs of *C. flexuosa* (*C. hirsuta* subgenome of *flexuosa*: F^h^ and *C. amara* subgenome of *flexuosa*: F^a^) and parents (*C. amara* genome: A; *C. hirsuta* genome: H) by PCA. Even though the same environmental factors were shown to be important for niche differentiation, the whole expression pattern was not clearly correlated with site, when we compared the sum expression of two subgenomes with the expression level of the parents (SI and Fig. S5). We then examined the expression level of the subgenomes separately. Two subgenomes were clearly divided into two large groups by PC1 with a 30% contribution rate, and the diploid genomes were located close to each daughter subgenome of the allopolyploid. PC2 roughly grouped each subgenome by date with 13% contribution rate, from early (downside) to late (upside) date (Fig. 3*B*). In the PC2 and PC3 plots, each study site formed loose group regardless of subgenomes (Fig. 3*C*). This result suggests that the whole genome expression pattern is strongly dominated by subgenome type as the primary factor, and subsequently by growth stage or environment. Similarly, the resemblance to the parental expression pattern was detected when the expression level of each homeolog of *C. flexuosa* was compared with that of the diploid parents by clustering analysis (Fig. S6).

### Differentially Expressed Genes and Gene Ontology Analysis

To examine how many genes are associated with habitat environment in the allopolyploid, we further analyzed the RNA-seq data of *C. flexuosa* to extract differentially expressed genes (DEGs) by considering the total expression of homeolog pairs. The number of genes used for this analysis is shown in Table S5*A*. We detected DEGs between site pairs among IR2, KT2 and KT5, on a given day using edgeR (44). The numbers of DEGs varied among combinations, with generally more genes detected between either of the two KT sites and IR1, and fewer within KT sites (Dataset S1 and Table S5*B*, see Fig. S7 for the summary). These results correspond to the geographical closeness of KT2 and KT5, and this trend persisted over time. Overall, more genes were differentially expressed among *C. flexuosa* on May 2 than other dates, and between KT2 and IR1 on any date, most probably reflecting the difference of the growth stages (Table S3).

We also detected DEGs between the allopolyploids and the cohabiting diploids in IR1 and KT5, and between diploids. The number of DEGs between H at IR1 and A at KT5 was the largest among all combinations, with 39% of analyzed genes detected as DEG (Table S5*B*, Fig. S7). To examine the types of genes involved in the response to various habitat environments, we conducted a Gene Ontology (GO) analysis on each DEGs set for *C. flexuosa* (Dataset S2, see SI for the detail). Many GO terms were shared among many of the nine site pair by date combinations; thus, we focused on the GO terms that appeared in five or more combinations (Table 6). Among them were GO:0009627 (systemic acquired resistance), many GO terms related to plant defense system, wounding, and oxidative stress, and GO:0009414 (response to water deprivation) were found in many combinations. These data suggest that biotic stress and water availability shape gene expression responses in the natural habitats we studied.

As the results of both habitat environment and gene expression analyses indicated that water was the key environmental factor for niche differentiation, we examined genes constituting the GO term GO:0009414 in our dataset. Few genes overlapped in the “within KT” and “between KT and IR” comparisons (Fig. S8*A*), and between different times (Fig. S8*B*), indicating that most genes were expressed at specific locations and times.

### Genes with Homeolog Expression Ratio Changes between Two Homeologs (HomeoRoq Genes) and GO Analysis

We used the HomeoRoq pipeline to examine the changes in the ratio (or bias) of homeolog expression levels between sites; i.e., we implemented a solid statistical test to identify homeologous pairs showing significant ratio changes by considering overdispersion (10) (Dataset S3). We examined approximately 20,000 genes (Table S5), which was nearly double the number of genes examined in the previous study (28). The number of homeolog pairs with significant expression ratio change (HomeoRoq genes) differed among site pair by date combinations and constituted 0.9%–3.3% of all genes (Table S5*A* and *B*). Similar to the case for the DEGs, the smallest number of genes was detected to be different between KT2 and KT5 on any date, and the biggest difference was found between IR1 and KT2 (Table S5*B*).

The GO analyses of the HomeoRoq genes identified 15 to 72 GO terms enriched in each site pair by date combination (Dataset S4, see SI for the detail). Among these GO terms, three terms, GO:0009414 (response to water deprivation), GO:0006979 (response to oxidative stress), and GO:0000302 (response to reactive oxidative species), were detected in the highest proportion, i.e., in six site combinations (Table S6). GO:0006833 (water transport) was found in five combinations.

We further examined the pattern of the gene expression in the GO:0009414 (response to water deprivation). Among 32 HomeoRoq genes in GO:0009414, 26 genes were also found as DEG (Table S7). Different homeologous pairs changed the ratio depending on sites and times (Fig. S9). To illustrate different patterns of expression changes, we visualized the expression levels in both allopolyploids and diploids of 24 genes in the GO:0009414 (Fig. S10), in which significant differences were detected both between allopolyploids (IR1 and KT5) and between two diploids (*C. hirsuta* in IR1 and *C. amara* in KT5). The gene CARHR207830, a homolog of the *Arabidopsis DREB2A* gene, was expressed higher in the drier IR1 site than in the wetter KT5 in all diploid and allopolyploid species, consistent with its protein function as a key transcriptional factor in drought response (45). In the allopolyploids in both sites, F^a^ showed three to five times higher expression than F^h^, suggesting *cis*-regulation, where diverged parental homeologs retain strong control (46, 47). The gene CHRHR100610 is a homolog of *Arabidopsis MBF1C*, a transcriptional coactivator induced by various stress including drought and heat and by plant hormone ABA and salicylic acid (48). F^h^ showed more than 10 times higher expression than F^a^ in both sites, suggesting strong *cis*-regulation. In the gene CARHR155190, a homolog of *Arabidopsis RDUF1* responsible for the response to a plant hormone ABA (49), F^a^ was lowly expressed at KT5, and thus was also detected as a gene with significant ratio change by HomeoRoq. These results suggest that there is no single dominant pattern of homeolog regulation in the natural environments and that the expression regulation depends on genes.

## Discussion

### The Allopolyploid *C. flexuosa* Combined its Parental Legacy in Fine-Scale Niche Acquisition

In this study, we documented the various environmental factors of the habitats of the allopolyploid and its diploid parents (Figs 1, 2, S2, and S3, Table 1). Together with previous growth experiments with water stress treatments (28), these data suggest the following information on niche differentiation of the two diploid parental species in *Cardamine* and the allopolyploid derived from them. First, the niches of the two diploid species are distinct. The habitats of the two species were separated in the PCA of various environmental factors (Fig. 1), among which soil water content contributed the most to niche differentiation after sky openness (Table 1, S2). The temporal data of water availability (Fig. 2) indicated that the habitats of *C. amara* and *C. hirsuta* were wet and dry, respectively. The fitness data of *C. hirsuta* (Table S2) supported its preference for dry habitat. These results are consistent with findings of the growth experiments detecting higher performance of *C. amara* under the submergence treatment and that of *C. hirsuta* under drought treatment (28). Second, the habitat of the allopolyploid was broad as indicated by the larger root-mean-square error along PC1 of Fig. 1, but the overlap with the parental species was limited at the intermediate range and thus excluded extreme environments (Fig. 1, Table 1). The broad range itself does not guarantee that the allopolyploid establishes its own niche. In the case of *C. flexuosa*, the temporal monitoring of habitat water availability revealed high fluctuation (Fig. 2), suggesting that the adaptation to a broad range of conditions can be advantageous in acquiring robustness in fluctuating habitat environments. Consistent with this observation, *C. flexuosa* grew similarly well under the submergence, drought, and the fluctuation of the two treatments in the laboratory experiment (28). Given these data, the allopolyploid *C. flexuosa* seems to have obtained a new distinct niche with fluctuating water availability.

By contrast with our results, broader niches for allopolyploids were not detected for *Primula* or *Glycine* using a modeling approach and climatic factors at a continental scale (50, 51). The inconsistency in the results from the present and previous studies might be because of differences in the taxa that were studied (varying in the genetic background and/or time since allopolyploidization), in spatial scale, in the environmental factors considered, and/or in the measurement methods (field-based or modeling approaches). Here, niche differentiation of the study species occurred at small geographical scales, often <1 km^2^ (e.g., *C. amara* in WBH 1 vs. *C. hirsuta* in WBH 2 and 3; Fig. S1), and we did not capture habitat differentiation at a larger geographic scale (IR, WBH, and KT). In addition, the occurrence of the study species was related to water content, C/N ratio, and sky openness at season start, but not to the other quantified environmental factors (Table 1) and not all factors were well correlated with one other (Table S8). Therefore, which factors are associated with niche differentiation depends on the factors included in the analysis. Given these limitations, our results underscore that empirical data on environmental factors contributing to species occurrence at a fine spatial scale are informative. This might lead to reevaluation of the proposed lack of niche differentiation in allopolyploids and diploid parents reported to date (19).

### The Subgenome-Wide Expression Pattern of *C. flexuosa* Resembles Respective Diploid Parents

Our results suggest that the gene expression pattern of each subgenome is still closer to each parent after the polyploidization 10,000–1 million years ago (33) because the PCA clearly separated subgenomes as the most dominant factor (Fig. 3*B*). The second influential factor in the PCA was the date, suggesting that the expression pattern of many of the genes are possibly attributed to the growth stage besides environment (Table S3). Compared with date, sites showed weaker effect on the expression pattern both in subgenome (Fig. 3*B* and *C*) and in total genome (Fig. S5). Similarly, clustering analysis grouped F^a^ with A separately from F^h^ with H (Fig. S6), regardless of habitat except for one F^a^ sample from KT5.

### Expression Pattern of Homeologs in the Allopolyploid *C. flexuosa* is Associated with the Key Factor in Niche Differentiation

Water availability was shown to be one of the key environmental factors in niche differentiation in *C. flexuosa* not only by ecological approach (Table 1), but also by RNA-seq analysis (Table S6). Indeed, water-related GO terms were detected in the majority of site pairs in both GO analyses based on DEG and on HomeoRoq genes (Table S6). One GO term (GO:0009414: response to water deprivation) was the same in our DEG and HomeoRoq analyses as in laboratory experiments manipulating water availability in *A. thaliana* (e.g., 52–54). Therefore, it seems that genes covered by GO:0009414 are commonly used in water responses in a range of environments in multiple species. These genes could have also been involved in the niche diversification between *C. amara* and *C. hirsuta*, and thus in the establishment of *C. flexuosa* in habitats of a wider range of water availability between progenitors. Diversification of the total expression levels as the result of differentially expressed homeologs may have conferred the plasticity for *C. flexuosa* to persist in a broad range of water availability.

GOs enriched among the field samples of *C. flexuosa* were similar to those enriched in *C. flexuosa* and the diploid parents in the previous laboratory experiment with submergence or drought treatments, despite differences in experimental (microarray vs. RNA-seq) and bioinformatic methods (28). Among water-related GOs, responses to abscisic acid, a plant hormone responsible for the drought response, was identified in both studies. In addition, response to reactive oxygen species (*C. hirsuta* by microarray in submergence) were enriched in both, which is consistent with the possibility of oxidative stress triggered by water-related stress. These similarities suggest that the same response mechanism as detected in the laboratory underlies niche differentiation in the field and further corroborate the importance of water-related responses in the *Cardamine* species in the field.

In this study, ecological and transcriptomic data indicated the same environmental factor for niche differentiation. The comparison between GO terms enriched by HomeoRoq and DEG analyses gives an insight into the underlying molecular mechanism. Many GO terms related to pathogen response were enriched in DEG analysis, but not in HomeoRoq analysis (Table S6). This suggests that the regulation of these genes is conserved in both parental genomes; therefore, they do not contribute to the niche differentiation, but rather reflect the difference of microbial flora in each site caused by water availability or other factors (55, 56). By contrast, water-related GO terms were enriched in both DEG and HomeoRoq analyses, suggesting that, concerning a key environmental factor for niche differentiation, an allopolyploid exhibits differential expression of homeologs in addition to its total expression level. More empirical data should allow us to examine whether this is commonly the case and whether we could specify the key environmental factor of allopolyploid niche differentiation based on expression patterns of DEG and HomeoRoq genes.

### The Allopolyploid Species Obtained a Generalist Niche by Transcriptomic Plasticity

The results of the present field study and previous laboratory experiments suggest that the allopolyploid *C. flexuosa* is a generalist both ecologically and transcriptomically, in that its habitat and gene expression cover a wide range. We found that *C. flexuosa* occurred in both wet and dry habitats. In the previous study, transcriptomic change of *C. flexuosa* was closer to that of *C. amara* by submerged treatment and to *C. hirsuta* by drought treatment (28). In the present study, the GOs of water- and ABA-related traits were enriched in DEGs and HomeoRoq genes among sites and times (Table S6), supporting the relevance of transcriptomic plasticity to water-related responses of *C. flexuosa*. These findings are consistent with the idea that allopolyploids obtained environmental robustness to adapt to a new generalist niche, and its major molecular mechanism is the transcriptomic plasticity by combining stress responses of their parental species (4, 12). The effect of complementarity is widespread in ecology, from reduced competition among individuals or enhanced productivity of species, to the stability of ecosystems (57–59). The combination of complementary genomes specialized in contrasting environments might result in a similar effect in allopolyploids possessing generalist properties.

A fundamental question on niche differentiation is how the species coexist (60). We propose three mutually nonexclusive reasons for the coexistence of the trio of *Cardamine* species. First, unlike parental diploids, *C. flexuosa* can thrive in fluctuating environments. Second, the diploids may prefer extreme environments, as suggested by higher fitness of *C. hirsuta* in a drier environment (Table S2), which may leave the intermediate environment open for *C. flexuosa*. Third, although *C. flexuosa* can occur in both wet and dry habitats, it may not be able to withstand extreme environments as much as the diploid parents due to attenuated response at the transcriptomic level. Indeed, the previous transcriptomic study showed that the degree of upregulation of stress response genes was in general attenuated in *C. flexuosa* compared with parental diploids (28). Similarly, in the allopolyploid *A. kamchatica*, the upregulation of dozens of key *cis*-regulated genes by metal treatment were reduced by up to half (12). Thus, it seems that becoming a generalist has a trade-off, i.e., its stress response is attenuated; therefore, it cannot live in the extremes of the environmental gradient. This concept of coexistence can be further examined by a manipulative field experiment where the three species are transplanted in a range of habitats from *C. amara* only to *C. hirsuta* only and evaluated for life history traits and transcriptomics.

The generalist niche and transcriptomic plasticity of an allopolyploid we documented here in *C. flexuosa* can be common to other allopolyploid species. For example, the genus *Cardamine* has been proposed to have experienced adaptive radiation along water availability gradients by recurrent allopolyploidization, and the anecdotally described habitat environments of allopolyploid species including *C. scutata* and *C. insueta* are intermediate and/or fluctuating (28), consistent with a generalist niche. At the transcriptomic level, genes involved in abiotic stress responses might be under *cis*-regulation not only in *C. flexuosa* (28), but also in other allopolyploid species, such as cotton, coffee, and *Arabidopsis* (5, 9, 13, 61, 62). Both in the present study (Table S5) and previous studies on Brassicaceae in the laboratory (10, 12), only a small portion (<4%) of all genes exhibited the homeolog expression ratio change. Taken together, a limited number of distinctively regulated environment-response homeologs may play a key role in the establishment of an allopolyploid in a generalist niche.

## Materials and Methods

Details not included here appear in SI.

### Study Sites

The study was conducted in seminatural and anthropogenic areas in and around Zurich, Switzerland (Fig. S1*A*, SI).

### Analyses on Habitat Environment

PCA and logistic regressions were conducted on 17 out of the 19 sites that had plants in both 2013 and 2014 (excluding IR9 and KT6). The data contained nine environmental factors: weight-based soil water content, soil C/N ratio, soil pH, NO_3_^−^ concentration in fresh soil, NH_4_^+^ concentration in fresh soil, NO_3_^−^ concentration in incubated soil, NH_4_^+^ concentration in incubated soil, sky openness at the beginning of the season, and sky openness at the end of the season. Further details are in SI.

### Transcriptomics Sample Preparation and RNA-seq

RNA was extracted from leaf tissue using an RNeasy Plant Mini Kit (Qiagen). Total RNA was used for the library synthesis using TruSeq Stranded mRNA Library Prep Kit (Illumina), and then sequenced using an Illumina HiSeq2500 at the Functional Genomics Center Zurich. On average 17.8 ± 4.1 million reads were obtained from 32 samples (Table S4). Because of its poor quality, one sample from IR1 on April 18 was excluded from the subsequent analyses. Sequence information has been deposited in the DNA Data Bank of Japan (DDBJ; http://www.ddbj.nig.ac.jp; RNA: DRA Accession ID: DRA006314, BioProject ID: PRJDB4989, BioSample ID: SAMD00097398-SAMD00097406, Assembly: DRA Accession ID: DRA006316, BioProject ID: PRJDB4989, BioSample ID: SAMD00098907).

### Reference-Guided Assembly of *C. amara* Genome and RNA-seq Read Quantification

We used the published genome sequence of *C. hirsuta* (34) and our *C. amara* genome sequence constructed by reference-guided de novo assembly based on *C. hirsuta* genome according to Lischer and Shimizu (63) (see SI for the detail). We targeted 29,458 homeolog pairs annotated in both A and H genomes. RNA-seq reads with low quality were trimmed with Trimmomatic (version 0.36) (64), then mapped to the *C. amara* and *C. hirsuta* genomes with STAR (version 2.5.3a) (65) to classify the origins with HomeoRoq (10, 66). The read number on each homeolog was counted with featureCounts (version 1.6.0) (67) and the proportion of the homeolog pairs was calculated (Table S4 and Fig. S4).

### DEG Analysis and Homeolog Expression Ratio Analysis

DEGs between two samples were identified by edgeR (version 3.22.2) (44) with a threshold of false discovery rate (FDR) <0.05. The sum of *C. hirsuta* origin reads, *C. amara* origin reads, and common (unclassified) reads by HomeoRoq was treated as the expression of one gene in *C. flexuosa* (see SI for the detail and Table S5*A* for detailed number of genes analyzed). The homologs of *A. thaliana* genes were identified by reciprocal best hit with BLASTN version 2.2.30^+^ (68).

HomeoRoq genes of the three sites on each of the three sampling dates were detected using HomeoRoq pipeline (10), with a cutoff threshold FDR <0.05. We applied FPKM >0.2 and ratio standard deviation among the sum of the biological replicates of the site pair <0.3 to all genes to filter out those with extremely low expression level and false positives, respectively. See SI for the detail and Table S5*A* for the detailed number of genes analyzed.

To characterize the type of genes being overrepresented among DEGs and HomeoRoq genes, we performed GO enrichment analyses among the nine site pair by date combinations and between cohabiting diploid and allopolyploid using the gene annotation according to *A. thaliana* (https://www.arabidopsis.org).

## Supporting information

DatasetS1_corr

SupportingInformation

## ACKNOWLEDGMENTS

The authors thank G. Halstead-Nussloch, M. Helling, A. Morishima, T. Paape, S. Röthlisberger, K.K.S. Ng, A. Tedder, M. Yamazaki, N. Zweifel, A. M. Suarez, and other members of the Shimizu Group, D. Schneebeli, S. Moor, P. Enz, B. Hirzel, D. Schlagenhauf, N. Kofmehl, H. Bär, M. Freund, P. Niklaus, R. Husi, A. Ferrari, T. Zwimpfer, M. Furler, B. Schmid, P. C. Joerg, R. Leiterer, D. Markulin, B. Kellenberger, J. Sugisaka, L. Koyama, K. Fukushima, and N. Sletvold for their advice and assistance. We also thank J. Kühn-Georgijevic at the Functional Genomics Center of Zurich for RNA-seq analysis; the Botanical Garden of the University of Zurich and Gemeinde Küsnacht-Tobel for permits for field studies; and Swisstopo for geographical data. This study was funded by the Swiss National Science Foundation to R.S.-I. and K.K.S.; by University Research Priority Programs, Evolution in Action of the University of Zurich to R.S.-I. and K.K.S.; by the Human Frontier Science Program to K.K.S., A.H., and Jun. S.; by the Japan Science and Technology Agency, Core Research for Evolutionary Science and Technology grant number JPMJCR16O3, Japan; and by KAKENHI grant numbers 16H06469 to K.K.S. and Jun. S., 18H04785 to K.K.S., and 26113709 and 16H01463 to M.M.K.

