## SupportingInformation for "Fine-scale ecological and transcriptomic data reveal niche differentiation of an allopolyploid from diploid parents in *Cardamine*"

**Study Species.** *Cardamine amara* L. (large bittercress) is a perennial widespread across most of central Europe (1–3). Populations of *C. amara* in lowland areas in Switzerland (<1,000 m above sea level) are reported to consist of diploids (2*n*=2*x*=16), and polyploid populations of subsp. *austriaca* have been observed at higher altitudes of the inner alpine valleys (3–5). Typically, *C. amara* grows to 10–60 cm in height and does not form a rosette (6). It propagates sexually and clonally (7). *Cardamine hirsuta* L. (hairy bittercress) is a diploid (2*n*=2*x*=16) native to Europe (1, 8, 9). It is 5–30 cm in height and forms a rosette (6). It is a winter annual and its seeds cannot tolerate submergence (10). *Cardamine flexuosa* With. (wavy bittercress) is an annual or perennial tetraploid (2*n*=4*x*=32) native to Europe (8), which originated from *C. hirsuta* and *C. amara* (1, 11). *Cardamine flexuosa* forms a rosette (12). Recently, plants that had been identified as *C. flexuosa* in eastern Asia turned out to belong to a distinct species, *C. occulta* (8, 13).

When not obvious, *C. flexuosa* and *C. hirsuta* were identified using morphological keys (12). The incidence of backcrossing is seemingly low, if at all, as we found none out of a total of 115 plants in the study sites screened for their ploidy level using flow cytometry (CyFlow Space, Partec, using Cystain UV precise P reagents). Specimens of the study species collected from the study sites have been donated to Zürich Herbaria, the herbarium of the University of Zurich and ETH Zurich, where they are registered with barcodes Z-000167685 to Z-000167691 for *C. hirsuta*, Z-000167692 to Z-000167697 for *C. flexuosa*, and Z-000167698 to Z-000167704 for *C. amara*.

**Study Sites.** The study was conducted in the seminatural and anthropogenic areas of Irchel, Wehrenbach, and Küsnacht-Tobel in and around Zurich, Switzerland (Fig. S1*A*). In 2013, we selected 18 sites (IR1–9, WBH1–3, KT1–5, 7) from the three areas (Fig. S1*B*–*D*). WBH1, WBH2, and WBH3 are separated by a hedge and a paved road. The composition of the species and the numbers of individuals of each species per site are summarized in Table S1. In 2014, the study was conducted in the same sites as in 2013, except for site KT6, which was newly added, and IR9, which was excluded because of its disappearance upon changed land use.

**Soil Properties.** To characterize the sites' soil properties, surface soil (0–5 cm depth, ca. 100 mL) beside each pot for moisture monitoring (see below) was sampled on May 14, 2013. The samples were kept cool during transportation to our laboratory at the University of Zurich, where they were passed through a 2-mm sieve. For water content and total content of carbon and nitrogen measurements, approximately 5 g of fresh sample was weighed before being dried at 105 °C for 48 h. The dried soil was weighed to calculate water content as the proportion of weight loss during drying. The soil was then milled, and total contents of carbon and nitrogen were quantified using a CHN analyzer (LECO Elemental Analyzer CHNS-932, LECO Instruments). For pH measurement, another subset of fresh sample was dried at 40 °C for two weeks, after which 5 g was dissolved in water to be measured with a handheld pH315i meter (WTW, Xylem Analytics Germany Sales). Inorganic nitrogen concentrations ($NO_3^-$ and $NH_4^+$) were quantified fresh and after a 4-week incubation (25 °C, RH 60%, dark, with MilliQ water supplied weekly to keep the original weight). Approximately 1.5 g of sample was extracted with 15 mL of 2M KCl and processed using a continuous flow analyzer (SKALAR San++).

**Soil Moisture Monitoring.** To monitor water availability of the study sites during the growing season, soil volumetric moisture was measured weekly from March to June in 2013 and 2014. The monitoring was done using an SM300 soil moisture sensor and an HH2 moisture meter (Delta-T Devices). These devices were designed to detect electrical conductivity in soil and are thus sensitive to soil composition, which was likely to vary among sites. To make measurements among sites comparable, we standardized the soil for moisture monitoring by burying unglazed pots (9 cm in height × 9 cm in diameter) filled with commercial soil (Floragard Floratorf, Floragard Vertriebs) to the ground level at each site. Three measurements were taken per pot and were averaged later. The pots were placed beside the study species, which were selected across the site. The number of pots per site ranged from three to eight, depending on the size of the site.

There was some irregularity in the monitoring in 2013: i.e., KT2 and KT3 were identified and added from the second and third readings, respectively. We lost some of the pots or the soil in them because of disturbances after heavy rain. In the latter cases, the soil was replaced, and the measurement was resumed from the following week to let the soil acclimate to the moisture content of its surroundings. Finally, the pots were removed as the plants dried up, resulting in the final census date being asynchronous among sites.

**Sky Openness.** To estimate sky openness at the sites, hemispheric photographs were taken at the pots used for moisture monitoring (see below) at each site using a digital camera (Nikon D90) with a circular fisheye lens (Sigma 4.5 mm f/2.8 EX DC circular fisheye) in 2014. To capture any seasonal trends, we took photographs twice in the season; in early spring and when the species were drying up. The first photographs were taken on March 24 or 26 at IR and WBH, and on March 22 at KT. The second photographs were taken for *C. hirsuta* on April 27, 28, or 29 at IR, WBH2 and WBH3, and for *C. flexuosa* and *C. amara* on June 18 at KT and WBH1. The photographs were manually converted to black and white images using the Fiji package of ImageJ software (https://fiji.sc/) before we calculated the proportion of the sky area using CanopOn2 (http://takenaka-akio.org/etc/canopon2/).

**Principal Component Analysis (PCA) on Habitat Environment.** The analysis was conducted on 17 sites where the species occurred in both 2013 and 2014 (see **Study Sites.** above). Out of the nine environmental factors measured, $NO_3^-$ concentration in fresh soil was excluded from analysis due to most of the values consisting of zeros and thus were not appropriate for PCA. The soil properties and sky openness of the sites were evaluated by PCA using software R-3.3.3 (14), with the “vegan” package (15). To examine the extent of variance of habitat environment of each species, we calculated the root-mean-square error for PC1 values. For each species for each year, we subtracted the mean PC1 value from PC1 value for each site, squared the outcome, summed the squared values to be divided by the number of sites, and then took the root of it.

**Modeling of Species Occurrence.** To examine the contributions of soil properties and light availability to the occurrence of each species in each year, we performed logistic regression analyses using PROC LOGISTIC in SAS version 9.3 (16). Out of the nine environmental factors measured, sky openness at the end of the season was excluded from the analysis due to its high collinearity with sky openness at the start of the season (Table S8). The site of occurrence of *C. flexuosa* was different in 2013 and 2014. Given this, we performed the analyses with following six factors as response variables: the occurrence (presence or absence) of *C. flexuosa* in 2013, *C. flexuosa* in 2014, *C. flexuosa* in both in 2013 and 2014, *C. flexuosa* either in 2013 or 2014, *C. amara*, and *C. hirsuta*. For each response variable, we ran 255 logistic regressions consisting of all possible combinations of the eight environmental factors (excluding the sky openness at the end of the season) as explanatory variables. Out of the 255 models examined, we identify the model with the lowest AIC value, which was regarded as the best model for the response variable. To circumvent separation issues, a Firth bias-corrected maximum likelihood adjustment was used for estimating regression coefficients. The significance of the explanatory variables in the best models was assessed using the log-likelihood ratio test (17).

**Fitness in Different Habitats for *C. hirsuta*.** To examine whether the study species might reproduce more at a particular soil moisture content and/or soil nutrient content, we analyzed the relationship between reproduction and weight-based soil water content and C/N ratio of the sites in 2013. Because most *C. amara* and *C. flexuosa* plants were lost from flooding, resulting in a low sample size, only the data for *C. hirsuta* were analyzed. At the beginning of the season, we marked and measured the length of the longest leaf of the rosette (as the initial size) of two plants beside the pot used for soil moisture monitoring. At the end of the season, we collected the plants to count the total numbers of fruits and estimate the mean number of seeds per fruit. By multiplying these two variables, we estimated the total number of seeds per plant as an index of fitness. Initial size and the total number of seeds were averaged between the two plants so that they could be comparable with the data on water content available per pot. In total, 39 samples were analyzed.

We regressed the total numbers of seeds against initial size, water content, and C/N ratio in linear and nonlinear multiple regression models using the statistical software R-3.3.3, with the "stats" package (14). Initial size was included in the model to control for its effect on response variable. Linear ($\beta$) and nonlinear ($\delta$) regression coefficients were extracted from respective models. Explanatory variables were standardized prior to analysis to enable direct comparison of results across traits. The total number of seeds was normalized by dividing each absolute value by the population mean (18). The variance inflation factor for initial size and water content did not exceed 2.1, indicating that collinearity was not a serious problem (19). Because the residuals from the multiple regressions had nonnormal distributions, statistical significance of regression coefficients was assessed by estimating 95% confidence intervals with bootstrapping, using the "boot" package (20). Data were resampled 5000 times and the confidence intervals were constructed based on saved coefficients from each run (21) with bias-correlated and accelerated methods (22).

**Tissue Sample Collection for Transcriptomics.** We selected three sampling sites in IR1, KT2, and KT5, where *C. flexuosa* coexisted with *C. hirsuta*, grew on its own, and coexisted with *C. amara*, respectively. The growth stage of each species at each sampling site and date of sampling is summarized in Table S3. Leaf tissue of *C. flexuosa* was collected from the same individuals on April 18, May 2, and May 16 in 2013, allowing for the exploration of temporal variation (23). The samples of the diploid parents were analyzed for only May 2, as *C. hirsuta* dried up by May 16 and *C. amara* was too small for sampling on April 18 and was lost to flooding after May 2. At each site, one individual beside a spot for soil moisture monitoring was marked at the beginning of the season. The samplings were conducted between 10:00 and 14:00 on sunny days to reduce any variation introduced by nonquantified stimuli, such as day/night cycles (23). Whenever possible, a fully developed and intact leaf before senescing was sampled from rosettes. On May 2, the leaf sample from one individual of *C. flexuosa* in KT2 was damaged. On May 16, a cauline leaf instead of a rosette leaf was sampled from one individual of *C. flexuosa* at IR1 because the rosette had dried up. The leaf samples were stored in a tube with RNAlater RNA Stabilization Reagent (Qiagen) and brought back to the laboratory. Once RNAlater infiltrated through leaves, the sample tubes were stored at –80 °C until further use.

**Reference-Guided *De Novo* Assembly and Annotation of *C. amara*.** For the genome sequences, one specific line of *C. amara* from Switzerland (lab ID: WEH ATS1) was clonally propagated and used for DNA extraction. We sequenced 368 million paired-end reads (3 independent libraries with different insertion sizes of 150–500 bp) and 548 million mate-pair reads (4 independent libraries with different insertion sizes from 3–15 kb). To annotate genes on *C. amara* genome, RNA samples from eight organs (anther, filament, sepal, petal, pistil, flower bud, young flower bud, and leaf) of another individual from Switzerland (lab ID: 3-19A) were sequenced. All sequence information used for assembly and annotation has been deposited in the DNA Data Bank of Japan (DDBJ; http://www.ddbj.nig.ac.jp) as DRA Accession ID: DRA006316, BioProject ID: PRJDB4989, BioSample ID: SAMD00098907.

All the *de novo* assemblies were performed according to a previous publication (24) using the ALLPATHS-LG version 51279 (25) tool. Final assembled scaffolds shorter than 1 kb were discarded. The assembled genome was annotated using the pipeline described in Briskine et al. (26) following the recommendations from the AUGUSTUS developers (27). Briefly, RNA-seq reads from eight tissues were aligned individually against the reference genome using STAR version 2.5.2a (28). The alignments were used to generate intron hints. In addition, we annotated repetitive elements with RepeatMasker version 4.0.6 (29) and RepBase Repeat Masker edition version 20140131 (30) derived *nonexonpart* hints from them.

Both sets of hints were used to generate the preliminary gene annotation with AUGUSTUS version 3.2 (31). The RNA-seq reads were aligned against exon–exon junction sequences extracted from the preliminary gene models using STAR. After removing spliced reads from the whole genome alignments, the remaining alignments were merged with exon–exon junction alignments. The merged alignments were then filtered to include only high-quality pairs, which were subsequently used to create a new set of intron hints for the final AUGUSTUS run.

To identify homologous genes of *Arabidopsis thaliana*, we applied a reciprocal best hit (RBH) method. We aligned gene region of the assembled genome onto *A. thaliana* gene region (TAIR10: https://www.arabidopsis.org) using BLASTN+ version 2.2.30 (32) with 1E-10 as a cutoff of E-value. The *A. thaliana* gene with the best BLAST result was defined as the homologous gene of the *C. hirsuta* gene.

**PCA by Total Genome of *C. flexuosa* and Parents.** We conducted PCA by the total expression level of two homeologous genes, at three time points (April 18, May 2, and May 16) at three sites with contrasting soil moisture levels (IR1, KT2, and KT5, Fig. 3*A*). We also included the cohabiting parents, *C. hirsuta* at IR1 and *C. amara* at KT5 on May 2. We calculated the fragments per kilobase per million (FPKM) of a total homeolog pair and filtered out the pairs (genes) with FPKM<1. This process left 23,065 homeolog pairs out of 29,458 known pairs, and we analyzed them by the same method as PCA by subgenomes (Fig. 3). The PCA was done using software R-3.5.0 (14), with *prcomp* in the “stats” package. By PC1 and PC2, whose contribution rates are 23% and 13%, each species was grouped (Fig. S5*A*). By PC1 and PC3 (contribution rate 12%), *C. flexuosa* was grouped by date, e.g., most probably reflecting the difference in growth stage (Fig. S5*B*). In contrast with the PCA with subgenomes (Fig. 3*C*), we could see less effect from sites in *C. flexuosa*. Overall, *C. flexuosa* samples are located closer to *C. amara* by PC1 and PC3, but closer to *C. hirsuta* by PC2.

**Clustering analysis of RNA-seq samples of *C. flexuosa* and parents.** The same homeolog expression data as PCA by subgenome (23,182 genes, Fig. 3) was used for clustering analysis to illustrate the relationship between each *C. flexuosa* homeolog ($F^a$ and $F^h$) and each parent (A and H) (Fig. S6). The analysis was done using *hclust* with Pearson's correlations in the "stats" package using R-3.5.0 (14).

**Differentially Expressed Gene and Gene Ontology Analysis.** Differentially expressed genes (DEGs) between site pairs on each sampling date were detected using edgeR with a false discovery rate (FDR) of <0.05 as the threshold value (33). In total, 29,458 genes for which homologs are known in the *A. thaliana* genome were subject to the analysis. For *C. flexuosa*, the sums of *C. hirsuta* origin reads, *C. amara* origin reads, and common (unclassified) reads (Fig. 1 in Ref. 34) were used for the analysis (Table S4), which was performed using the SUSHI framework (35). The number of detected DEGs of each site pair by date combination is summarized in Table S5, Fig. S7, and Dataset S1.

At the next step, a Gene Ontology (GO) enrichment analysis of the biological process was conducted using topGO version 2.32.0 with the elim algorithm at a significance level of $p = 0.05$ (36) for each site pair on each sampling date to examine which gene categories are overrepresented in each habitat. For this GO analysis, only the genes remained after RBH with *A. thaliana* genes (29,458 genes) were targeted as parent population. The results of all comparisons can be found as Dataset 2. GO terms with <10 or >500 annotated genes were excluded from interpretation.

By the comparison of two parents, *C. amara* in KT5 and *C. hirsuta* at IR1 on May 2, 39% of the total genes were detected as DEGs, suggesting the largest difference among the comparisons shown in Fig. S7. Many of the detected GO terms were related to the cell wall, pathogenic response, and oxidative stress (Dataset 2). In addition, GO:0009414 (response to water deprivation) was also detected (Dataset 2). These findings could be attributed to the difference of the growth stage of plants and the difference in habitat environment, in addition to the difference between species. While *C. hirsuta* already started senescing at IR1, *C. amara* was not flowering yet and the leaf was still under expansion on May 2 (Table S3). In addition, the difference in soil moisture level in the surrounding may shape the difference in pathogenic composition or abundance (37, 38).

We also compared *C. flexuosa* and the cohabiting parent at KT5 and IR1, respectively. In PCA plots, *Cardamine flexuosa* samples were separated from samples of both parents, which were separated from each other (Fig. S5). These results indicate that the expression pattern at whole genome level diverges between an allopolyploid and its diploid parents and between two diploids, and may reflect the difference among subgenomes and the difference in the growth stage (Table S3).

Compared with the difference between allopolyploid and the cohabiting diploid, the number of DEGs among *C. flexuosa* samples was generally smaller (Fig. S7 and Table S5). The number of *C. flexuosa* DEGs between IR1 and KT2 was larger than that between KT2 and KT5 or between IR1 and KT5. These results may reflect the shorter geographical distance between KT2 and KT5 than other site pairs (Fig. S1). It may also reflect that the growth stage of *C. flexuosa* was similar between IR1 and KT5 but not between the other site pairs (Table S3), suggesting a substantial influence of the growth stage, besides habitat environment, on expression patterns. Among nine site pair by date combinations, GO terms related to defense response to fungus and bacteria were listed in majority of them (Dataset S2). The pathogenic flora might have differed among habitats, because water availability can affect the activity and flora of pathogens (37, 38).

**Genes with Homeolog Expression Ratio Change (HomeoRoq genes) and GO Analysis.** The homeolog expression ratio *H* is expressed by:

$$H(g_a, g_h) = \frac{g_a}{g_a + g_h}, \quad (1)$$

where $g_a, g_h$ are the expressions of homeologous genes derived from *C. amara* and *C. hirsuta*, respectively. Changes in the homeolog expression ratios among sites on each sampling date was examined using the HomeoRoq pipeline (34), with FDR <0.05 as the threshold for significance. In all, 18,855 genes were subject to the analysis. These were homologs of *A. thaliana* genes identified by RBH with BLASTN version 2.2.30$^+$ (32) and were a subset of 24,670 genes that fulfilled the criteria for the following filtering step. Among the 29,458 genes for which homologs existed in *A. thaliana*, we only analyzed genes whose standard deviation among the sum of the biological replicates of the site pair of concern was <0.3 to remove false positives and whose FPKM was >0.2 to exclude extremely poorly expressed genes. The number of HomeoRoq genes are summarized in Table S5 and Fig. S7.

The number of homeolog pairs detected by HomeoRoq was generally smaller than DEGs in the same comparison pairs (Table S5, see Fig. S7 for summary), corresponding to only 0.9%–3.3% of the total homeolog pairs, suggesting that the expression of most homeolog pairs are similarly regulated even at a different site and date. GO analysis of biological process was conducted for the detected gene in each combination by targeting only the genes for which *A. thaliana* homologs were identified with RBH (29,458 genes). We used topGO version 2.32.0 with the elim algorithm at a significance level of $p = 0.05$ (36). GO terms with <10 or >500 annotated genes were excluded from interpretation. The results of all comparison can be found as Dataset 4.

**Overlap of Genes in GO:0009414 between Different Site Pairs and between Dates.** For each of DEG and HomeoRoq genes, we examined how many genes overlapped between different site pairs and between dates, using R-4.1.0, with the package “eulerr” (39). The results are shown as Venn diagram in Fig. S8 (DEG) and Fig. S9 (HomeoRoq genes).

**SI Figure Legends**

**Fig. S1** Locations, compositions, and photos of representative sites of the study species. (*A*) Locations of the study areas Irchel (IR), Wehrenbach (WBH), and Küsnacht-Tobel (KT) in and around Zurich, Switzerland. (*B*) Locations of sites at IR. (*C*) Locations of sites at WBH. (*D*) Locations of sites at KT. (*E*) The *C. hirsuta* site. (*F*) The *C. flexuosa* site. (*G*) The *C. amara* sites. In (*B–D*), different colors of points indicate different species composition: red, *C. hirsuta* only; orange, *C. hirsuta* and *C. flexuosa*; green, *C. flexuosa* only; blue, *C. flexuosa* and *C. amara*; and black, *C. amara* only. The left and right halves of the circles represent the years 2013 and 2014, respectively, for the sites where the species composition differed between years. In (*E–F*), an overview (top) and a hemispheric photo used for light availability analysis (bottom) are shown with date and site name.

**Fig. S2** Soil properties from 18 sites in 2013. (*A*) water content (*B*) C/N ratio (*C*) $NO_3^-$ concentration in fresh soil (*D*) $NH_4^+$ concentration in fresh soil (*E*) $NO_3^-$ concentration in incubated soil (*F*) $NH_4^+$ concentration in incubated soil (*G*) pH. The points and background are colored according to Figs. S1 and 1, respectively. In (*C*), the level was not detectable except for IR1-3.

**Fig. S3** Sky openness from 18 sites measured at the season's start (around Julian calendar day 80) and season's end (around Julian calendar day 120 for *C. hirsuta* sites and 170 for *C. flexuosa* and *C. amara* sites, respectively) in 2014. The points and lines are colored as in Fig. S1.

**Fig. S4** The proportion of *C. hirsuta* origin and *C. amara* origin reads in RNA-seq samples. Each row constitutes one sample. Species, ploidy level, sampling site, and sampling date for each sampling are shown. The third replicate of *C. flexuosa* on April 18 was not included due to the low quality of the sequence.

**Fig. S5** The result of PCA on the expression level of 23,065 genes. The total expression of a homeolog pair was treated as one gene expression for *C. flexuosa*. Circles in black and red indicate samples of the diploid parents *C. amara* and *C. hirsuta*, respectively. (*A*) The colors of points correspond to dates: light brown, April 18; brown, May 2; dark brown, May 16. (*B*) The points are colored as in Fig. S1.

**Fig. S6** The result of a clustering analysis based on $\log_{10}$FPKM calculated for 23,182 genes in *C. amara* (A), *C. hirsuta* (H), *C. amara* homeolog of *C. flexuosa* ($F^a$), and *C. hirsuta* homeolog of *C. flexuosa* ($F^h$) on May 2.

**Fig. S7** A summary of the number of HomeoRoq genes of 23,182 genes analyzed and DEGs of 23,065 genes analyzed of the study species on May 2.

**Fig. S8** *(A)* Proportional Venn diagrams for the number of genes in category GO:0009414 (response to water deprivation) overlapping between site pairs on each date, based on the GO enrichment analysis for DEGs between site pairs. *(B)* Proportional Venn diagrams for the number of genes in GO:0009414 (response to water deprivation) overlapping between dates for each site pair, based on the GO enrichment analysis for DEGs between site pairs. We analyzed 113 genes.

**Fig. S9** *(A)* Proportional Venn diagrams for the number of genes in the category GO:0009414 (response to water deprivation) overlapping between site pairs on each date, based on the GO enrichment analysis on the HomeoRoq genes. *(B)* Proportional Venn diagrams for the number of genes in the category GO:0009414 (response to water deprivation) overlapping between dates for each site pair, based on the same GO enrichment analysis as (A). We analyzed 32 genes.

**Fig. S10** The expression level of 24 genes in GO:0009414 (response to water deprivation) for which significant differences were detected both between allopolyploids (*C. flexuosa* in IR1 and in KT5) and between diploids (*C. hirsuta* in IR1 and *C. amara* in KT5). The headings show the *C. hirsuta* gene ID with the *A. thaliana* homolog ID and gene name in parenthesis. H and A indicate *C. hirsuta* and *C. amara* genes, whereas $F^h$ and $F^a$ indicate *C. hirsuta* and *C. amara* homeologs of *C. flexuosa*, respectively.

**Dataset S1** Excel file of DEGs (FDR <0.05 in edgeR). Each sheet is for a site pair on one date. The sheet names consist of the species (A: *C. amara*, F: *C. flexuosa*, or H: *C. hirsuta*), site (IR1, KT2, or KT5), and date (April 18, May 2, or May 16). For example, the sheet A_KT5_May2_vs_H_IR1_May2 contains DEGs between *C. amara* at KT5 on May 2 and *C. hirsuta* at IR1 on May 2. Each sheet consists of the following columns: *C. hirsuta* gene ID, ATID: *A. thaliana* gene ID, Symbols: symbols for the gene, Full gene name: full name of the gene, log2-fold-change: fold change of the gene expression level when the sample before 'vs' of the sheet name is used as the control, FDR: false discovery rate.

**Dataset S2** Excel file of GO lists for DEG. Only GOs with >10 and <500 annotated genes are presented. Each sheet is for a site pair on one date. The sheets are named the same way as they are in Dataset S1. Each sheet consists of the following columns: GO ID, Term: GO term, Annotated: the number of genes annotated in the target population (see SI, page 9) for the GO term, Significant: the number of significant genes detected for the GO term in the study dataset, Expected: the expected number of significant genes for the GO term, *p*: *p*-value according to topGO elim.

**Dataset S3** Excel file of genes that changed the homeolog expression ratio (FDR <0.05 in HomeoRoq). Each sheet is for a site pair on one date. The sheets are named the same way as they are in Dataset S1. Each sheet consists of the following columns: *C. hirsuta* gene ID, ATID: *A. thaliana* gene ID, Symbols: symbols for the gene, Full gene name: full name of the gene, FDR: false discovery rate, ratioSD: standard deviation among the samples from the two groups of concern.

**Dataset S4** Excel file of GO lists for genes that changed the homeolog expression ratio. Only GOs with >10 and <500 annotated genes are presented. Each sheet is for a site pair on one date. The sheets are named the same way as they are in Dataset S1. Each sheet consists of the following columns: GO ID, Term: GO term, Annotated: the number of genes annotated in the target population (see SI, page 9) for the GO term, Significant: the number of significant genes detected for the GO term in the study dataset, Expected: the expected number of significant genes for the GO term, *p*: *p*-value according to topGO elim.

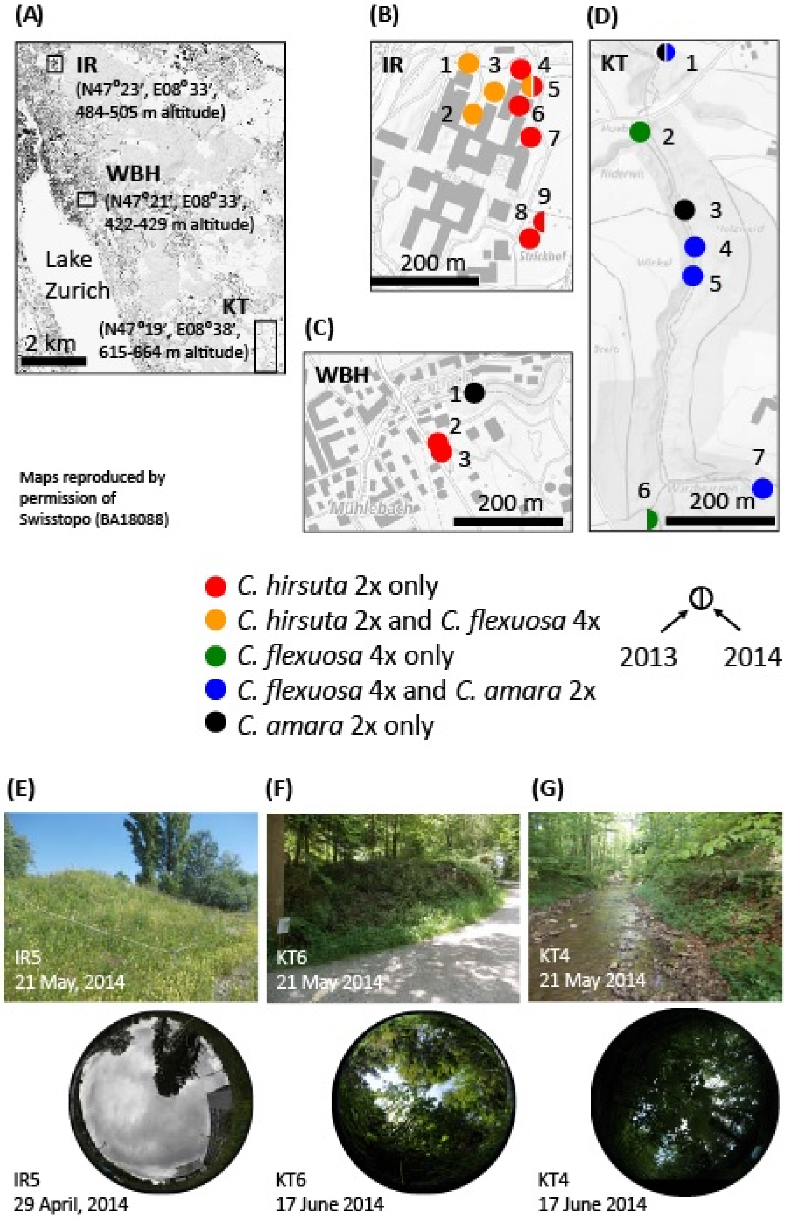


**Fig. S1.** Locations, compositions, and photos of representative sites of the study species. *(A)* Locations of the study areas Irchel (IR), Wehrenbach (WBH), and Küsnacht-Tobel (KT) in and around Zurich, Switzerland. (*B*) Locations of sites at IR. (*C*) Locations of sites at WBH. (*D*) Locations of sites at KT. (*E*) The *C. hirsuta* site. (*F*) The *C. flexuosa* site. (*G*) The *C. amara* sites. In (*B–D*), different colors of points indicate different species composition: red, *C. hirsuta* only; orange, *C. hirsuta* and *C. flexuosa*; green, *C. flexuosa* only; blue, *C. flexuosa* and *C. amara*; and black, *C. amara* only. The left and right halves of the circles represent the years 2013 and 2014, respectively, for the sites where the species composition differed between years. In (*E–F*), an overview (top) and a hemispheric photo used for light availability analysis (bottom) are shown with date and site name.

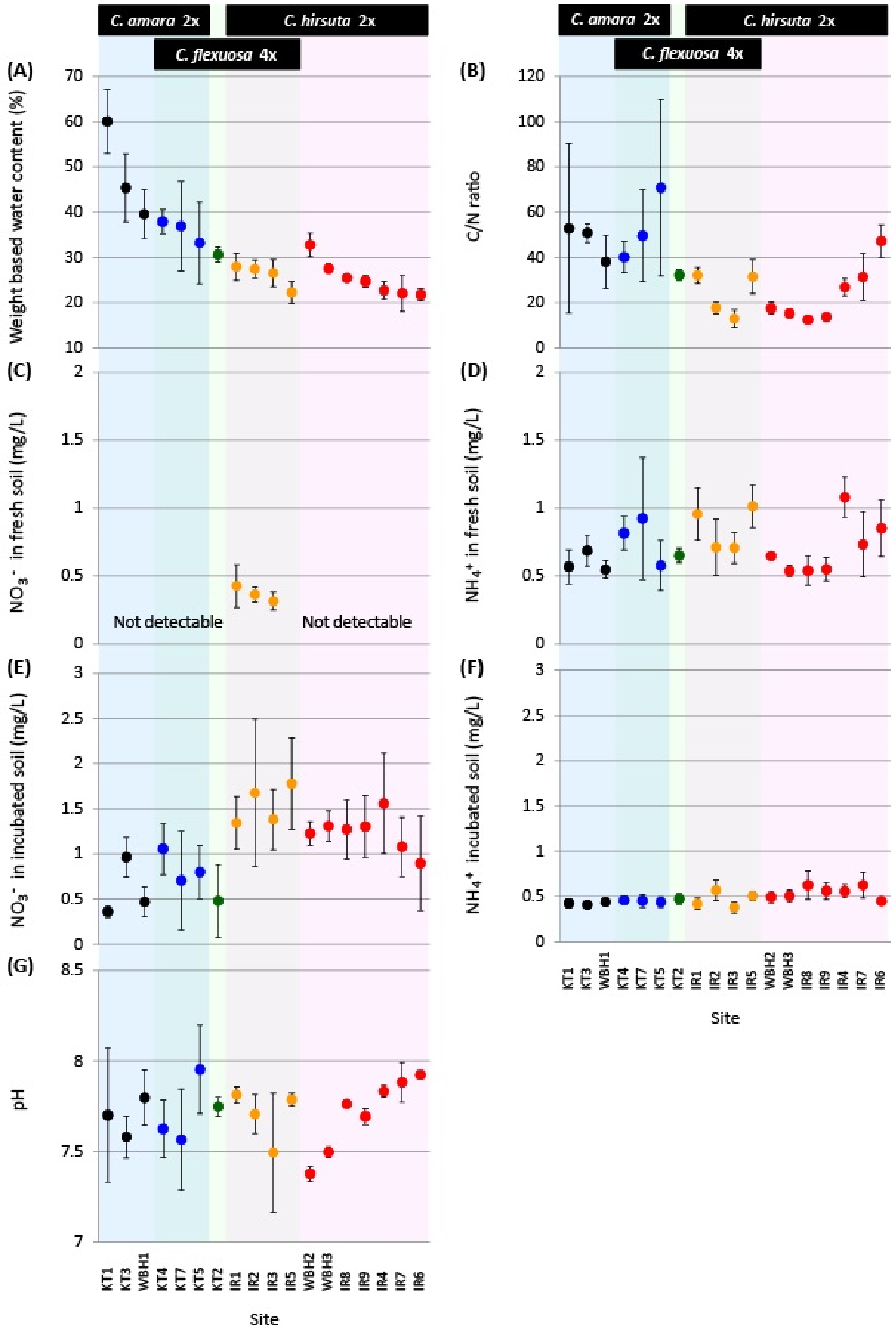


**Fig. S2.** Soil properties from 18 sites in 2013. (*A*) water content (*B*) C/N ratio (*C*) $NO_3^-$ concentration in fresh soil (*D*) $NH_4^+$ concentration in fresh soil (*E*) $NO_3^-$ concentration in incubated soil (*F*) $NH_4^+$ concentration in incubated soil (*G*) pH. The points and background are colored according to SI Appendix, Figs. S1 and 1, respectively. In (*C*), the level was not detectable except for IR1-3.

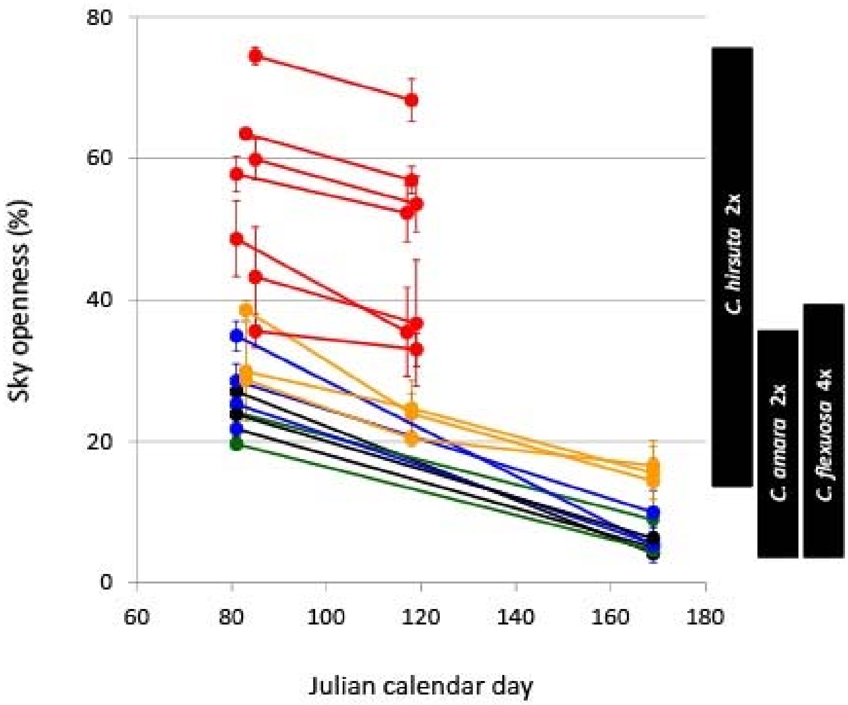


**Fig. S3.** Sky openness from 18 sites measured at the season's start (around Julian calendar day 80) and season's end (around Julian calendar day 120 for *C. hirsuta* sites and 170 for *C. flexuosa* and *C. amara* sites, respectively) in 2014. The points and lines are colored as in SI Appendix, Fig. S1.

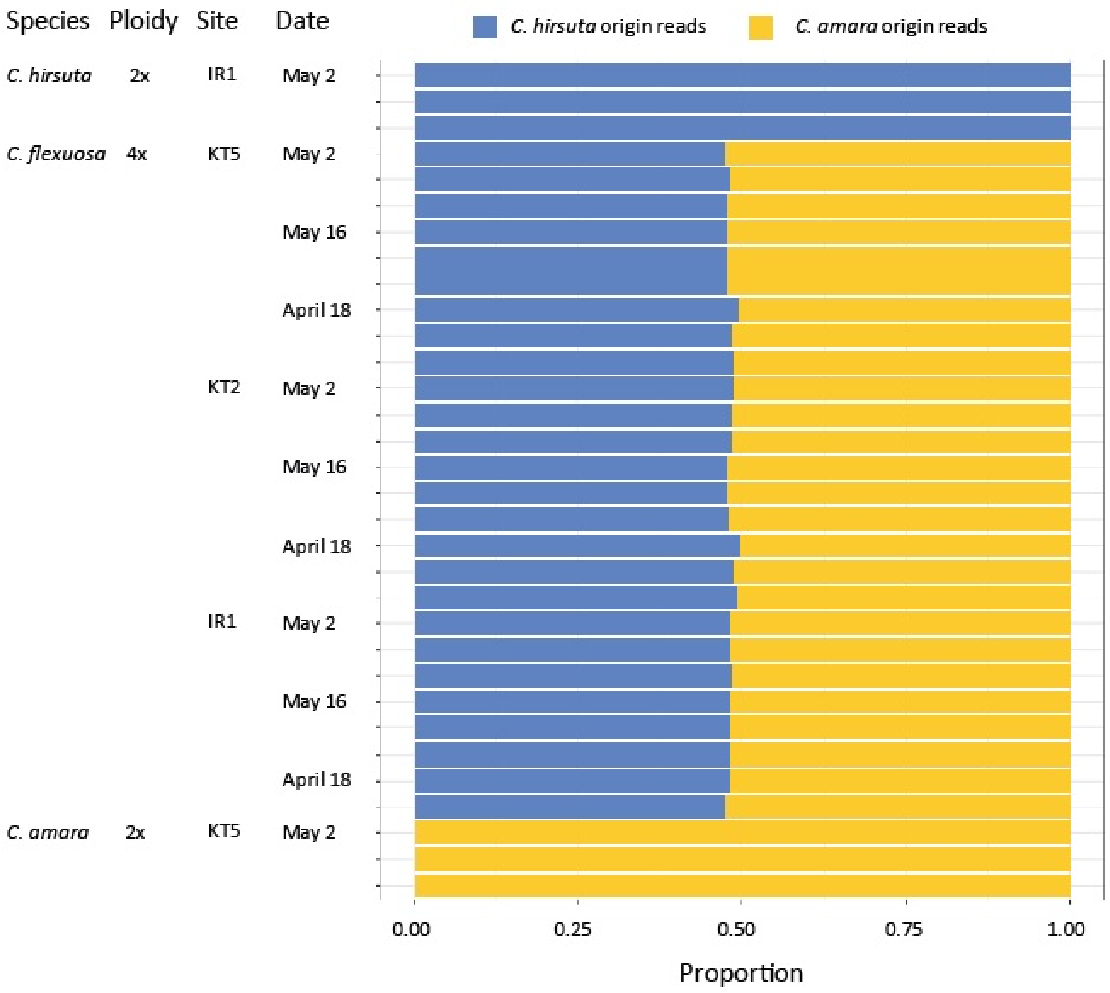


**Fig. S4.** The proportion of *C. hirsuta* origin and *C. amara* origin reads in RNA-seq samples. Each row constitutes one sample. Species, ploidy level, sampling site, and sampling date for each sampling are shown. The third replicate of *C. flexuosa* on April 18 was not included due to the low quality of the sequence.

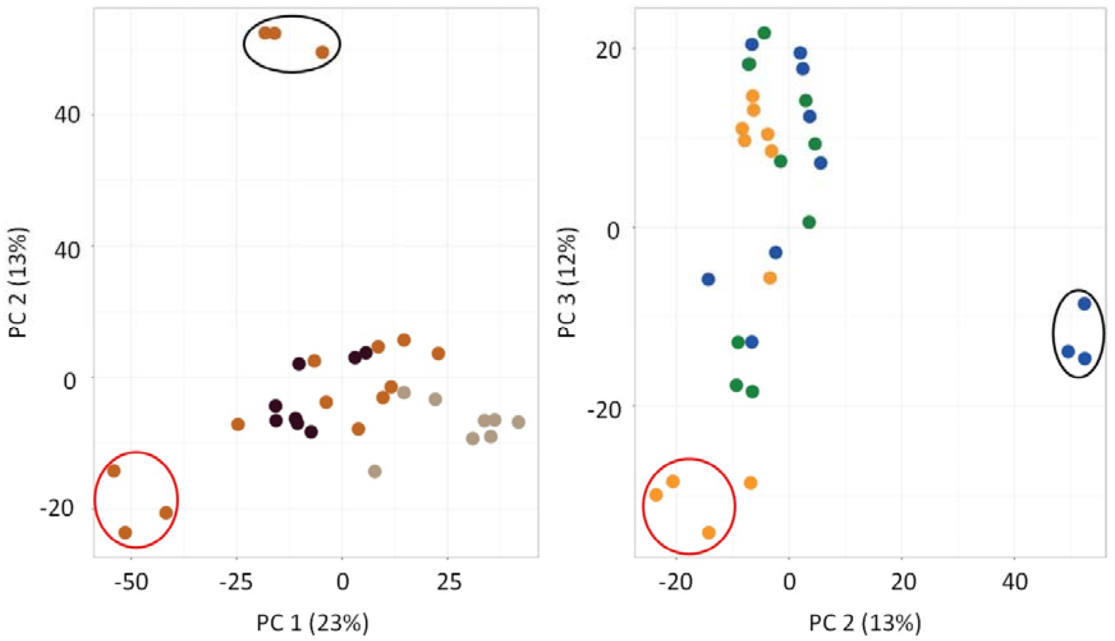


**Fig. S5.** The result of PCA on the expression level of 23,065 genes. The total expression of a homeolog pair was treated as one gene expression for *C. flexuosa*. Circles in black and red indicate samples of the diploid parents *C. amara* and *C. hirsuta*, respectively. (*A*) The colors of points correspond to dates: light brown, April 18; brown, May 2; dark brown, May 16. (*B*) The points are colored as in SI Appendix, Fig. S1.

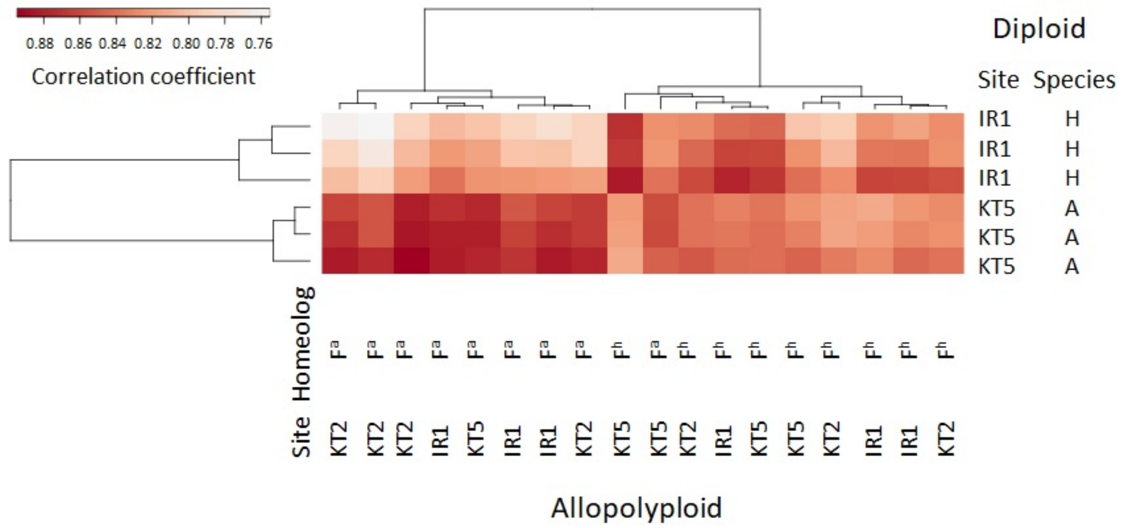


**Fig. S6.** The result of a clustering analysis based on $\log_{10}$FPKM calculated for 23,182 genes in *C. amara* (A), *C. hirsuta* (H), *C. amara* homeolog of *C. flexuosa* ($F^a$), and *C. hirsuta* homeolog of *C. flexuosa* ($F^h$) on May 2.

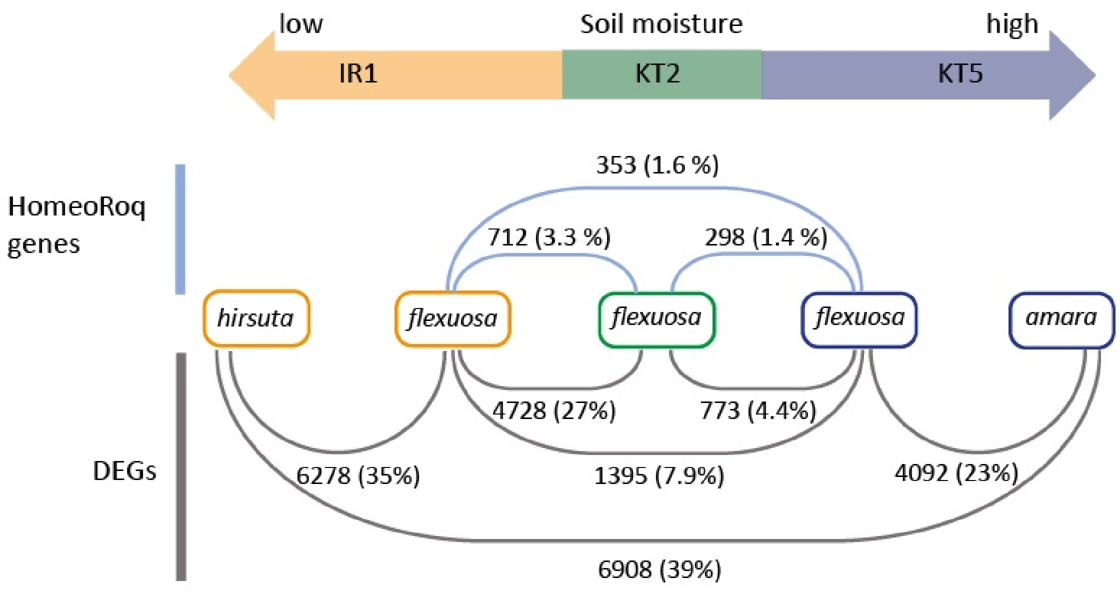


**Fig. S7.** A summary of the number of HomeoRoq genes of 23,182 genes analyzed and DEGs of 23,065 genes analyzed of the study species on May 2.

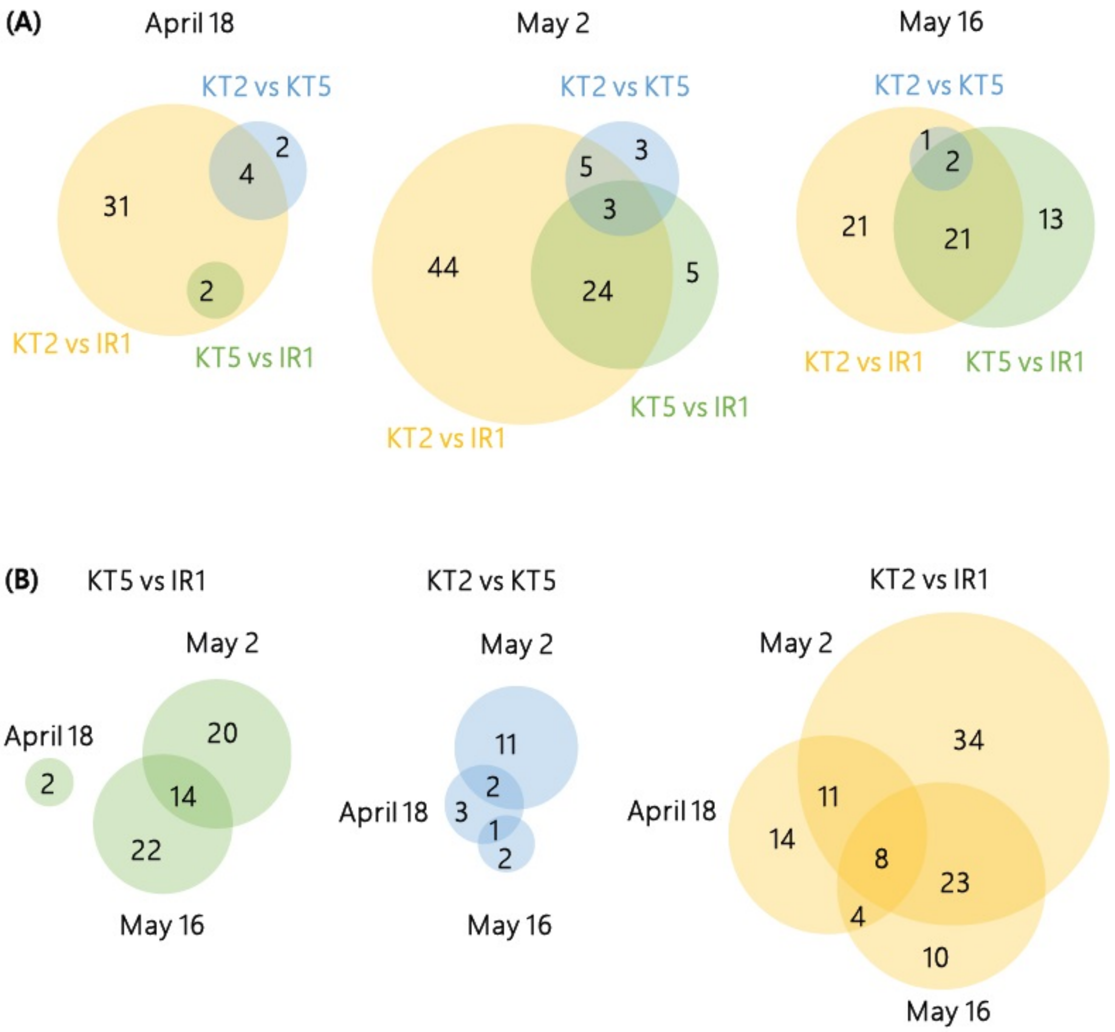


**Fig. S8.** *(A)* Proportional Venn diagrams for the number of genes in category GO:0009414 (response to water deprivation) overlapping between site pairs on each date of *C. flexuosa*, based on the GO enrichment analysis for DEGs between site pairs. *(B)* Proportional Venn diagrams for the number of genes in GO:0009414 (response to water deprivation) overlapping between dates for each site pair of *C. flexuosa*, based on the GO enrichment analysis for DEGs between site pairs. We analyzed 113 genes.

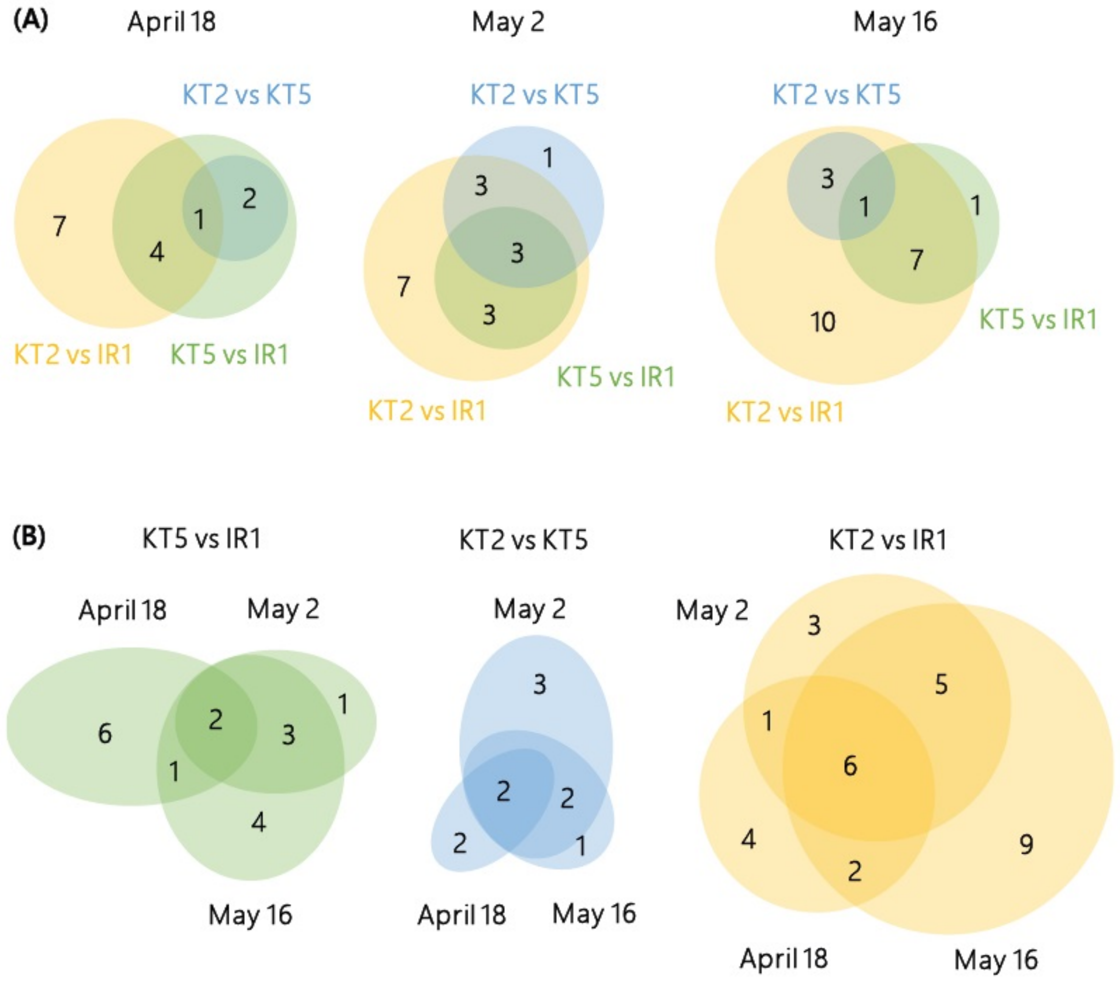


**Fig. S9.** *(A)* Proportional Venn diagrams for the number of genes in the category GO:0009414 (response to water deprivation) overlapping between site pairs on each date of *C. flexuosa*, based on the GO enrichment analysis on the HomeoRoq genes. *(B)* Proportional Venn diagrams for the number of genes in the category GO:0009414 (response to water deprivation) overlapping between dates for each site pair of *C. flexuosa*, based on the same GO enrichment analysis as (A). We analyzed 32 genes.

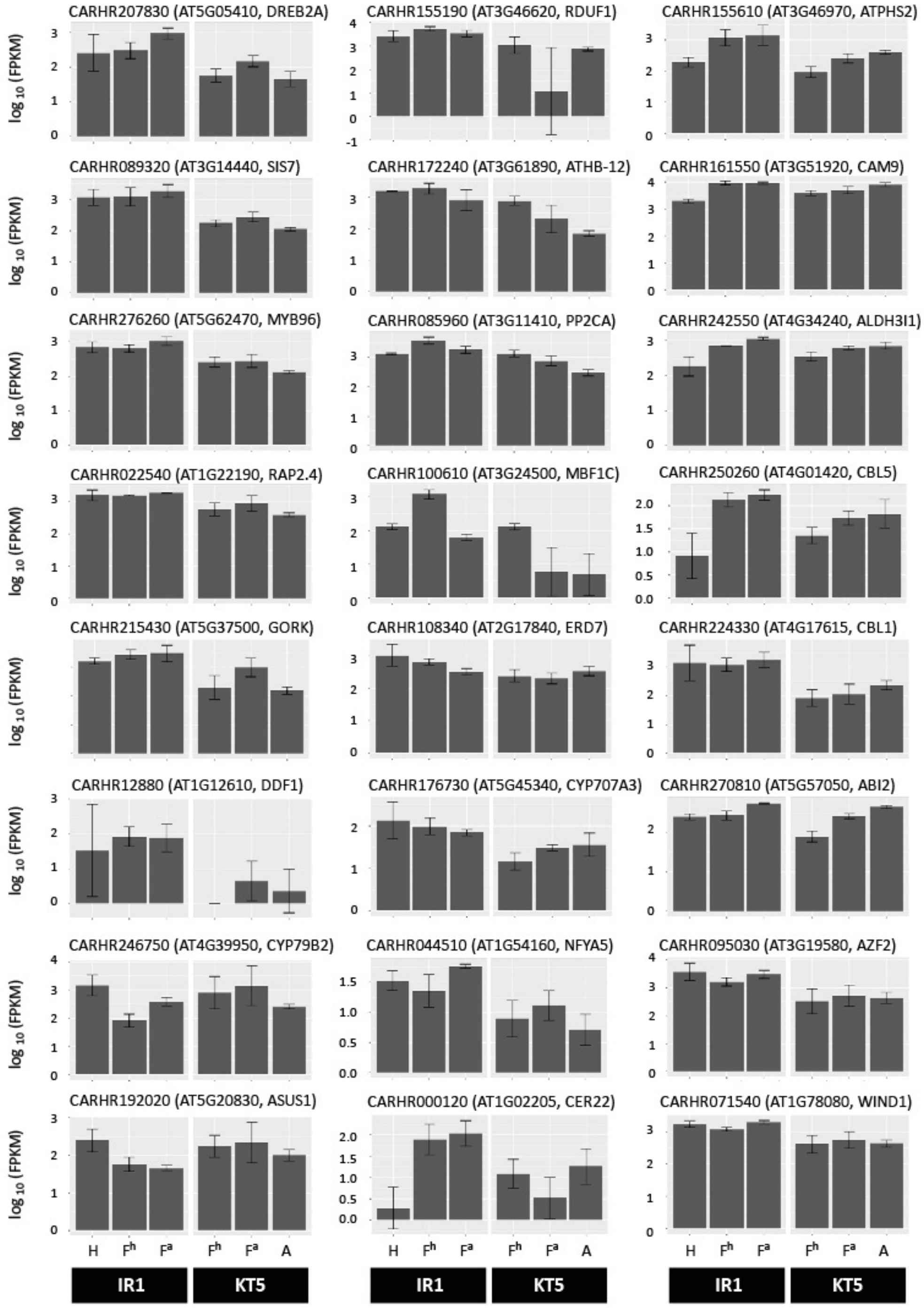


**Fig. S10.** The expression level of 24 genes in GO:0009414 (response to water deprivation) for which significant differences were detected both between allopolyploids (*C. flexuosa* in IR1 and in KT5) and between diploids (*C. hirsuta* in IR1 and *C. amara* in KT5). The headings show the *C. hirsuta* gene ID with the *A. thaliana* homolog ID and gene name in parenthesis. H and A indicate *C. hirsuta* and *C. amara* genes, whereas $F^h$ and $F^a$ indicate *C. hirsuta* and *C. amara* homeologs of *C. flexuosa*, respectively.

**Table S1.** The name of the study areas, the name and species composition (Type) of the study sites, and the number of individuals of the study species in 2013 and 2014. The number of individuals was recorded during March 4 to April 11 in 2013 and on April 10 in 2014. In Type, a = *C. amara* (2x), f = *C. flexuosa* (4x), h = *C. hirsuta* (2x). NAs indicate data not available. KT6 and KT9 were excluded from PCA on habitat environments.

| | | 2013 | | | | 2014 | | | |
|---|---|---|---|---|---|---|---|---|---|
| Area | Site | *C. amara* | *C. flexuosa* | *C. hirsuta* | Type | *C. amara* | *C. flexuosa* | *C. hirsuta* | Type |
| Irchel | IR1 | 0 | 25 | 160 | hf | 0 | 8 | 52 | hf |
| | IR2 | 0 | 6824 | 61 | hf | 0 | 332 | 64 | hf |
| | IR3 | 0 | 20 | 20 | hf | 0 | 11 | 6 | hf |
| | IR4 | 0 | 0 | 162 | h | 0 | 0 | 123 | h |
| | IR5 | 0 | 16 | 189 | hf | 0 | 0 | 411 | h |
| | IR6 | 0 | 0 | 170 | h | 0 | 0 | 146 | h |
| | IR7 | 0 | 0 | 280 | h | 0 | 0 | 64 | h |
| | IR8 | 0 | 0 | 46 | h | 0 | 0 | 425 | h |
| | IR9 | 0 | 0 | 276 | h | NA | NA | NA | NA |
| Wehrenbach | WBH1 | 750 | 0 | 0 | a | 700 | 2 | 0 | a |
| | WBH2 | 0 | 0 | 132 | h | 0 | 0 | 28 | h |
| | WBH3 | 0 | 0 | 205 | h | 0 | 1 | 161 | h |
| Küsnacht-Tobel | KT1 | 223 | 0 | 0 | a | 1190 | 25 | 0 | fa |
| | KT2 | 0 | 211 | 0 | f | 0 | 126 | 0 | f |
| | KT3 | 430 | 1 | 0 | a | 800 | 1 | 0 | A |
| | KT4 | 247 | 60 | 0 | fa | 640 | 28 | 0 | fa |
| | KT5 | 210 | 61 | 0 | fa | 305 | 74 | 0 | fa |
| | KT6 | NA | NA | NA | NA | 0 | 169 | 0 | f |
| | KT7 | 276 | 10 | 0 | fa | 640 | 39 | 0 | fa |

**Table S2.** Effects of initial size and water content on the total number of seeds per plant in *Cardamine hirsuta* (2x) in 2013 ($N = 39$) analysed with linear and non-linear multiple regressions. Partial regression coefficients with 95% confidence intervals in parenthesis are given. Statistically significant regression coefficients are in bold.

| Type of regression | $R^2$ | Initial size | Water content | CN ratio | Water content × CN ratio |
|---|---|---|---|---|---|
| Linear regression coefficient ($\beta$) | 0.32 | -0.078 (-0.377, 0.472) | **-0.461** (-1.102, -0.155) | **-1.025** (-1.739, -0.483) | - |
| Non-linear regression coefficient ($\delta$) | 0.56 | -0.052 (-0.226, 0.212) | **0.401** (0.049, 1.049) | **0.896** (0.399, 1.575) | **1.156** (0.403, 2.293) |

**Table S3.** The developmental stage at sampling of the three individuals of *Cardamine hirsuta*, *C. flexuosa*, and *C. amara* in IR1, KT2, and KT5 in 2013. B = Bolting, F = Flowering, S = Silique

| | IR1 | | | | | | KT2 | | | KT5 | | | | | |
|---|---|---|---|---|---|---|---|---|---|---|---|---|---|---|---|
| | *C. hirsuta* (2x) | | | *C. flexuosa* (4x) | | | *C. flexuosa* (4x) | | | *C. flexuosa* (4x) | | | *C. amara* (2x) | | |
| Date | 1 | 2 | 3 | 1 | 2 | 3 | 1 | 2 | 3 | 1 | 2 | 3 | 1 | 2 | 3 |
| April 18 | | | | B | B | B | B | B | B | B | B | B | | | |
| May 2 | F + S | F + S | F + S | F | F | F | B | B | B | F | F | F | B | B | B |
| May 16 | | | | S | F + S | F + S | F + S | F + S | F + S | F + S | F + S | F + S | | | |

**Table S4** The numbers of reads mapped by HomeoRoq (H-origin: mapped to *Cardamine hirsuta* (2x); A-origin: mapped to *C. amara* (2x)) and the number of expressed homeologs (FPKM>1) in parenthesis shown for each individual of *C. hirsuta*, *C. flexuosa* (4x), and *C. amara*. The second sample of *C. flexuosa* from IR1 on April 18 had a poor mapping quality and therefore excluded from the analyses.

| Species | Site | Date | H-origin | A-origin | Unclassified | Note |
|---|---|---|---|---|---|---|
| *C. hirsuta* | IR1 | May 2 | 6,831,864 (16,090) | 57,890 ( - ) | 124,787 | |
| | IR1 | May 2 | 9,928,515 (18,231) | 94,351 ( - ) | 244,915 | |
| | IR1 | May 2 | 9,944,365 (18,134) | 101,159 ( - ) | 217,253 | |
| *C. flexuosa* | IR1 | April 18 | 2,604,299 (15,721) | 2,990,519 (16,099) | 303,454 | |
| | IR1 | April 18 | 900,736 ( 5,429) | 1,050,553 ( 5,719) | 122,160 | Not analyzed |
| | IR1 | April 18 | 2,920,867 (13,388) | 3,357,210 (13,829) | 425,009 | |
| | IR1 | May 2 | 3,400,601 (17,416) | 3,766,561 (17,942) | 379,901 | |
| | IR1 | May 2 | 3,932,227 (17,025) | 4,431,741 (17,634) | 442,516 | |
| | IR1 | May 2 | 4,387,576 (17,390) | 4,872,114 (17,975) | 482,412 | |
| | IR1 | May 16 | 4,312,741 (17,009) | 4,889,803 (17,584) | 448,572 | |
| | IR1 | May 16 | 2,412,540 (16,760) | 2,722,960 (17,223) | 258,058 | |
| | IR1 | May 16 | 2,714,887 (16,567) | 3,073,875 (17,013) | 298,709 | |
| | KT2 | April 18 | 3,622,938 (16,842) | 3,965,621 (17,365) | 414,247 | |
| | KT2 | April 18 | 3,513,535 (16,255) | 3,887,057 (16,768) | 410,985 | |
| | KT2 | April 18 | 3,872,097 (16,912) | 4,154,868 (17,332) | 450,913 | |
| | KT2 | May 2 | 4,822,438 (17,238) | 5,430,549 (17,705) | 564,642 | |
| | KT2 | May 2 | 3,397,687 (17,221) | 3,814,394 (17,718) | 401,235 | |
| | KT2 | May 2 | 2,909,748 (16,867) | 3,235,626 (17,445) | 349,844 | |
| | KT2 | May 16 | 4,830,756 (17,617) | 5,557,942 (18,117) | 518,674 | |
| | KT2 | May 16 | 3,548,682 (17,113) | 4,150,121 (17,613) | 390,246 | |
| | KT2 | May 16 | 3,285,931 (17,107) | 3,828,325 (17,606) | 361,974 | |
| | KT5 | April 18 | 3,267,687 (16,380) | 3,579,829 (16,749) | 371,415 | |
| | KT5 | April 18 | 3,324,636 (17,052) | 3,672,647 (17,550) | 356,643 | |
| | KT5 | April 18 | 3,448,567 (17,244) | 3,724,853 (17,713) | 388,189 | |
| | KT5 | May 2 | 3,337,598 (17,300) | 3,855,335 (17,832) | 390,480 | |
| | KT5 | May 2 | 3,737,000 (17,201) | 4,249,544 (17,760) | 453,096 | |
| | KT5 | May 2 | 4,598,723 (17,340) | 5,279,825 (17,818) | 504,789 | |
| | KT5 | May 16 | 3,300,477 (16,875) | 3,815,878 (17,382) | 380,569 | |
| | KT5 | May 16 | 3,217,628 (16,909) | 3,713,061 (17,442) | 357,668 | |
| | KT5 | May 16 | 4,130,500 (17,317) | 4,716,904 (17,771) | 436,618 | |
| *C. amara* | KT5 | May 2 | 82,204 ( - ) | 6,702,228 (17,584) | 234,400 | |
| | KT5 | May 2 | 101,324 ( - ) | 8,804,071 (17,815) | 261,919 | |
| | KT5 | May 2 | 41,160 ( - ) | 4,169,433 (16,781) | 91,593 | |

**Table S5.** *(A)* The numbers of genes that were examined in DEG and homeolog expression ratio change (HomeoRoqs) analyses b[illegible] sites on three sampling dates in *Cardamine flexuosa* (4x), *C. amara* (2x), and *C. hirsuta* (2x). The numbers in parentheses indicate genes for which *A. thaliana* homolog exist according to Reciprocal Best Hit (RBH). *(B)* The numbers of genes that turned out to b[illegible] DEG and homeolog expression ratio change (HomeoRoqs) analyses between pairs of sites on three sampling dates in *C. flexuosa* ( (2x), and *C. hirsuta* (2x). The numbers in parentheses indicate the number of genes for which *A. thaliana* homolog exist according

| Table | Analysis | Site pair | April 18 | May 2 | May 16 |
|---|---|---|---|---|---|
| (A) | DEG | *C. flexuosa* KT5 vs. *C. flexuosa* IR1 | 17,832 (15,328) | 17,564 (15,121) | 17,357 (14,927) |
| | | *C. flexuosa* KT2 vs. *C. flexuosa* KT5 | 17,757 (15,306) | 17,563 (15,106) | 17,376 (14,934) |
| | | *C. flexuosa* KT2 vs. *C. flexuosa* IR1 | 17,624 (15,176) | 17,316 (14,944) | 17,481 (15,010) |
| | | *C. flexuosa* IR1 vs. *C. hirsuta* IR1 | | 17,866 (15,213) | |
| | | *C. flexuosa* KT5 vs. *C. amara* KT5 | | 17,862 (15,186) | |
| | | *C. hirsuta* IR1 vs. *C. amara* KT5 | | 17,873 (15,101) | |
| | HomeoRoqs | *C. flexuosa* KT5 vs. *C. flexuosa* IR1 | 20,784 (16,926) | 21,507 (17,243) | 21,155 (17,049) |
| | | *C. flexuosa* KT2 vs. *C. flexuosa* KT5 | 21,213 (17,138) | 21,285 (17,074) | 21,317 (17,120) |
| | | *C. flexuosa* KT2 vs. *C. flexuosa* IR1 | 20,502 (16,755) | 21,378 (17,159) | 21,357 (17,164) |
| (B) | DEG | *C. flexuosa* KT5 vs. *C. flexuosa* IR1 | 225 (169) | 1,395 (1,139) | 1,292 (1,046) |
| | | *C. flexuosa* KT2 vs. *C. flexuosa* KT5 | 343 (259) | 773 (631) | 329 (232) |
| | | *C. flexuosa* KT2 vs. *C. flexuosa* IR1 | 2,231 (1,790) | 4,728 (3,992) | 3,470 (2,809) |
| | | *C. flexuosa* IR1 vs. *C. hirsuta* IR1 | | 6,278 (5,040) | |
| | | *C. flexuosa* KT5 vs. *C. amara* KT5 | | 4,092 (3,138) | |
| | | *C. hirsuta* IR1 vs. *C. amara* KT5 | | 6,908 (5,449) | |
| | HomeoRoqs | *C. flexuosa* KT5 vs. *C. flexuosa* IR1 | 417 (340) | 353 (272) | 269 (210) |
| | | *C. flexuosa* KT2 vs. *C. flexuosa* KT5 | 195 (144) | 298 (227) | 215 (157) |
| | | *C. flexuosa* KT2 vs. *C. flexuosa* IR1 | 642 (508) | 712 (565) | 648 (515) |

**Table S6.** Summary of GOs that were detected for the majority of site by time combinations for genes whose homeolog expression ratio significantly changed (HomeoRoq) and DEG in the allopolyploid *Cardamine flexuosa* (4x). Site pairs for which a given GO was enriched is indicated by x.

| Data type | Number of combinations | GO ID | GO term | Number of annotated genes in the GO | April 18 | | | May 2 | | | May 16 | | |
|---|---|---|---|---|---|---|---|---|---|---|---|---|---|
| | | | | | KT2 vs IR1 | KT2 vs KT5 | KT5 vs IR1 | KT2 vs IR1 | KT2 vs KT5 | KT5 vs IR1 | KT2 vs IR1 | KT2 vs KT5 | KT5 vs IR1 |
| HomeoRoq | 6 | GO:0009414 | response to water deprivation | 318 | x | | x | x | x | | x | | x |
| | | GO:0006979 | response to oxidative stress | 397 | x | | | x | x | x | x | | x |
| | | GO:0000302 | response to reactive oxygen species | 151 | x | | | x | x | x | x | | x |
| | 5 | GO:0006833 | water transport | 18 | | x | | x | | | x | x | x |
| DEG | 7 | GO:0009627 | systemic acquired resistance | 61 | x | | x | x | | x | x | x | x |
| | 6 | GO:0009408 | response to heat | 179 | x | | | x | | x | x | x | x |
| | | GO:0009751 | response to salicylic acid | 180 | x | | | x | | x | x | x | x |
| | | GO:0050832 | defense response to fungus | 237 | x | | | x | | x | x | x | x |
| | | GO:0010200 | response to chitin | 125 | x | | | x | | x | x | x | x |
| | | GO:0045490 | pectin catabolic process | 76 | x | | | x | | x | x | x | x |
| | | GO:0042545 | cell wall modification | 137 | x | | | x | x | | x | x | x |
| | 5 | GO:0009414 | response to water deprivation | 318 | x | | | x | x | x | | | x |
| | | GO:0009611 | response to wounding | 188 | x | | | x | | x | | x | x |
| | | GO:0009737 | response to abscisic acid | 493 | x | | | x | x | x | | | x |
| | | GO:0009753 | response to jasmonic acid | 206 | x | x | | | | x | x | | x |
| | | GO:0042542 | response to hydrogen peroxide | 59 | x | | | x | | x | x | | x |
| | | GO:0009644 | response to high light intensity | 72 | x | | | x | | x | x | | x |
| | | GO:0042493 | response to drug | 490 | x | | | x | | | x | x | x |
| | | GO:0042335 | cuticle development | 30 | x | | x | x | | x | x | | |
| | | GO:0042742 | defense response to bacterium | 340 | | | | x | | x | x | x | x |

**Table S7.** List of *Cardamine hirsuta* genes and description of their *Arabidopsis thaliana* homologs belonging to GO:0009414 identified in the GO analysis on DEG that changed the homeolog expression ratio between at least one pair of sites on at least one date. The symbols and descriptions were acquired from TAIR website: https://www.arabidopsis.org/

| *C. hirsuta* gene ID | *A. thaliana* gene ID | Symbols | Description |
|---|---|---|---|
| CARHR000120 | AT1G02205 | CER22, CER1 | aldehyde catabolic process, alkane biosynthetic process, cuticle development, defense response to bacterium, defense response to fungus, response to water deprivation, wax biosynthetic process |
| CARHR027240 | AT1G27730 | ZAT10, STZ | multicellular organism growth, negative regulation of transcription, DNA-templated, photoprotection, photosynthesis, response to abscisic acid, response to chitin, response to cold, response to high light intensity, response to oxidative stress, response to salt stress, response to water deprivation, response to wounding |
| CARHR043110 | AT1G52890 | NAC019, ANAC019 | multicellular organism development, positive regulation of transcription, DNA-templated, response to water deprivation |
| CARHR073790 | AT1G80710 | DRS1 | cellular response to DNA damage stimulus, regulation of DNA damage checkpoint, response to abscisic acid, response to water deprivation |
| CARHR080150 | AT3G05700 | | positive regulation of transcription, DNA-templated, response to abscisic acid, response to salt stress, response to water deprivation |
| CARHR086990 | AT3G12490 | ATCYSB, ATCYS6, CYSB | defense response, hyperosmotic response, response to cold, response to oxidative stress, response to water deprivation |
| CARHR105210 | AT2G14610 | ATPR1, PR1, PR 1 | defense response, response to vitamin B1, response to water deprivation, systemic acquired resistance |
| CARHR155190 | AT3G46620 | RDUF1, AtRDUF1 | abscisic acid-activated signaling pathway, protein autoubiquitination, protein ubiquitination, response to abscisic acid, response to chitin, response to water deprivation |
| CARHR155610 | AT3G46970 | ATPHS2, PHS2 | glycogen catabolic process, response to cadmium ion, response to water deprivation |
| CARHR156890 | AT3G48170 | ALDH10A9, BADH | oxidation-reduction process, response to abscisic acid, response to water deprivation |
| CARHR224330 | AT4G17615 | CBL1, ATCBL1, SCABP5 | abscisic acid-activated signaling pathway, calcium-mediated signaling, pollen tube growth, response to cold, response to osmotic stress, response to salt stress, response to water deprivation, stomatal movement |
| CARHR232360 | AT4G24960 | HVA22D, ATHVA22D | flower development, hyperosmotic salinity response, negative regulation of autophagy, pollen development, response to abscisic acid, response to cold, response to water deprivation |
| CARHR004640 | AT1G05180 | AXR1 | response to cytokinin |

| | | | |
|---|---|---|---|
| CARHR034560 | AT1G35720 | ANNAT1, ATOXY5, OXY5, ANN1, AtANN1 | calcium ion transmembrane transport, phloem sucrose unloading, potassium ion export across plasma membrane, primary root development, response to abscisic acid, response to cadmium ion, response to cold, response to heat, response to osmotic stress, response to oxidative stress, response to salt stress, response to water deprivation |
| CARHR154040 | AT3G29320 | PHS1 | glycogen catabolic process, response to temperature stimulus, response to water deprivation |
| CARHR163320 | AT3G53420 | PIP2;1, AtPIP2;1, PIP2A, PIP2 | hydrogen peroxide transmembrane transport, response to abscisic acid, response to water deprivation, transmembrane transport, water transport |
| CARHR171770 | AT3G61430 | ATPIP1, PIP1A, PIP1, PIP1;1 | response to water deprivation, transmembrane transport, water transport |
| CARHR172240 | AT3G61890 | ATHB-12, ATHB12, HB-12 | multicellular organism development, positive regulation of transcription, DNA-templated, regulation of transcription, DNA-templated, response to abscisic acid, response to osmotic stress, response to salt stress, response to virus, response to water deprivation |
| CARHR062690 | AT1G69700 | ATHVA22C, HVA22C | hyperosmotic salinity response, response to abscisic acid, response to water deprivation |
| CARHR071540 | AT1G78080 | AtWIND1, WIND1, RAP2.4 | cellular response to salt stress, cytokinin-activated signaling pathway, ethylene-activated signaling pathway, red or far-red light signaling pathway, regulation of cell differentiation, regulation of transcription, DNA-templated, response to cold, response to light stimulus, response to osmotic stress, response to salt stress, response to water deprivation, response to wounding |
| CARHR093850 | AT3G18490 | ASPG1 | protein catabolic process, proteolysis, response to abscisic acid, response to water deprivation, systemic acquired resistance |
| CARHR098970 | AT3G23050 | IAA7, AXR2 | auxin-activated signaling pathway, gravitropism, regulation of transcription, DNA-templated, response to auxin, response to jasmonic acid, response to water deprivation, response to wounding |
| CARHR131680 | AT2G37170 | PIP2;2, PIP2B | response to abscisic acid, response to water deprivation, transmembrane transport, water transport |
| CARHR246750 | AT4G39950 | CYP79B2 | camalexin biosynthetic process, defense response, defense response by callose deposition in cell wall, defense response to bacterium, defense response to oomycetes, indoleacetic acid biosynthetic process, induced systemic resistance, oxidation-reduction process, response to bacterium, response to insect, response to water deprivation, tryptophan catabolic process |
| CARHR261630 | AT5G40390 | RS5, SIP1 | mannitol biosynthetic process, raffinose family oligosaccharide biosynthetic process, response to abscisic acid, response to oxidative stress, response to water deprivation, sucrose biosynthetic process |
| CARHR280630 | AT5G67030 | LOS6, NPQ2, ZEP, ABA1, ATABA1, ATZEP, IBS3 | abscisic acid biosynthetic process, oxidation-reduction process, response to heat, response to osmotic stress, response to red light, response to water deprivation, sugar mediated signaling pathway, xanthophyll biosynthetic process |

**Table S8.** Pearson coefficients from pairwise correlations between environmental variables.

| | Water content | CN ratio | pH | $NO_3$ | $NH_4$ | $NO_3$ incubated | $NH_4$ incubated | Sky openness at season start | Sky openness at season end |
|---|---|---|---|---|---|---|---|---|---|
| Water content | - | 0.507 * | -0.267 | -0.208 | -0.397 | -0.662 ** | -0.501 * | -0.425 | -0.603 * |
| CN ratio | | - | 0.423 | -0.349 | 0.029 | -0.612 ** | -0.467 | -0.543 * | -0.570 * |
| pH | | | - | -0.061 | 0.207 | -0.108 | 0.230 | -0.047 | 0.019 |
| $NO_3$ | | | | - | 0.181 | 0.430 | -0.174 | -0.174 | -0.205 |
| $NH_4$ | | | | | - | 0.459 | 0.011 | 0.200 | 0.216 |
| $NO_3$ incubated | | | | | | - | 0.411 | 0.583 * | 0.612 ** |
| $NH_4$ incubated | | | | | | | - | 0.618 ** | 0.685 ** |
| Sky openness at season start | | | | | | | | - | 0.959 *** |
| Sky openness at season end | | | | | | | | | - |

*$P < 0.05$, **$P < 0.01$, *** $P < 0.001$
